# Estimating essential phenotypic and molecular traits from integrative biodiversity data

**DOI:** 10.1101/2024.04.02.587699

**Authors:** Kristian Peters, Jörg Ziegler, Steffen Neumann

**Affiliations:** German Centre for Integrative Biodiversity Research (iDiv) Halle-Jena-Leipzig, Puschstraße 4, 04103 Leipzig, Germany; Institute of Biology/Geobotany and Botanical Garden, Martin Luther University Halle-Wittenberg, Am Kirchtor 1, 06108 Halle (Saale), Germany; Computational Plant Biochemistry, Leibniz Institute of Plant Biochemistry, Weinberg 3, 06120 Halle (Saale), Germany; Bioinformatics and Systems Biology, Justus Liebig University Giessen, Heinrich-Buff-Ring 58, 35392 Gießen, Germany; Metabolomics Facility, Program Center MetaCom, Leibniz Institute of Plant Biochemistry, Weinberg 3, 06120 Halle (Saale), Germany

## Abstract

In the context of biodiversity, only few functional traits and mechanisms are known from underrepresented groups such as mosses (bryophytes). Here, we use 16 field samples of complex thallose liverworts (order Marchantiales) collected from biological soil crusts as reference data for the reusable computational framework iESTIMATE that integrates and extracts phenotypic and molecular traits; and estimates Essential Molecular Variables (EMV). Our reference data involves (1) bioimaging, (2) metabolomics, and (3) DNA marker sequencing. These data are used to demonstrate the systematic and standardized extraction of phenotypic and molecular traits. To demonstrate the reusability of our framework, we propose naming schemes, apply Random Forest to estimate EMVs, phylogenetic dendrograms and partitioning around medoids to connect evolutionary relationships with ecological hypotheses and to document knowledge gains across domains. With this work we want to encourage the combined assessment, reuse and integration of phenotypic and molecular traits into functional ecology, biodiversity and related disciplines.

## Introduction

Vascular land plants (tracheophytes) have evolved a remarkable diversity of form and function, yet comparably few traits have been described in non-vascular plants such as bryophytes^1^. For vascular plants, functional traits can be reduced to two main axes representing either morphological characters such as sizes of plants and their parts, or on the second dimension the leaf economics spectrum which directly relates to molecular functioning and metabolic processes^2,3^. However, for the majority of bryophytes no analogous categories are available^4^. Despite the fact that bryophytes share a common ancestor with all tracheophytes, 450 million years of diversification led to a considerable macroevolutionary divergence and morphological disparity – in which mechanisms, patterns and functional processes are still largely unresolved^5,6^.

Bryophytes are composed of mosses, liverworts and hornworts. Unlike vascular plants, they have dominant gametophytes, lack seeds, shoot apical meristems, true roots and vascular tissues^7^. They further lack well-differentiated organs that protect them from environmental exposures and pathogens. As a result, classic phenotypic traits such as those involving plant sizes or discernible plant parts^2^ are often difficult to assess. By contrast, molecular traits have recently been shown to explain additional axes of specialisation in tracheophytes^8^ supporting the hypothesis that bryophytes interact with their biotic and abiotic environment predominantly at the molecular scale via small molecules^9,10^. In order to resolve this hypothesis, experimental data and methodological frameworks are needed to systematically assess traits in bryophytes^1^.

Here, we demonstrate reference data of 16 field samples of complex thallose liverworts (order Marchantiales) collected from different biological soil crust communities in Southern Sweden and Germany (Figure 1, Table 1). Thallose liverworts are of considerable interest for gaining insights into land plant evolution, historical radiation events, molecular mechanisms and morphological innovations^5,7^. They significantly contribute to ecosystem services and impact biogeochemical cycles at a global level^11^. As the mechanisms are still largely unknown, determining the essential traits comprising phenotypic and molecular levels can greatly help to assess their functional roles in ecosystems^11^.

**Figure 1.**
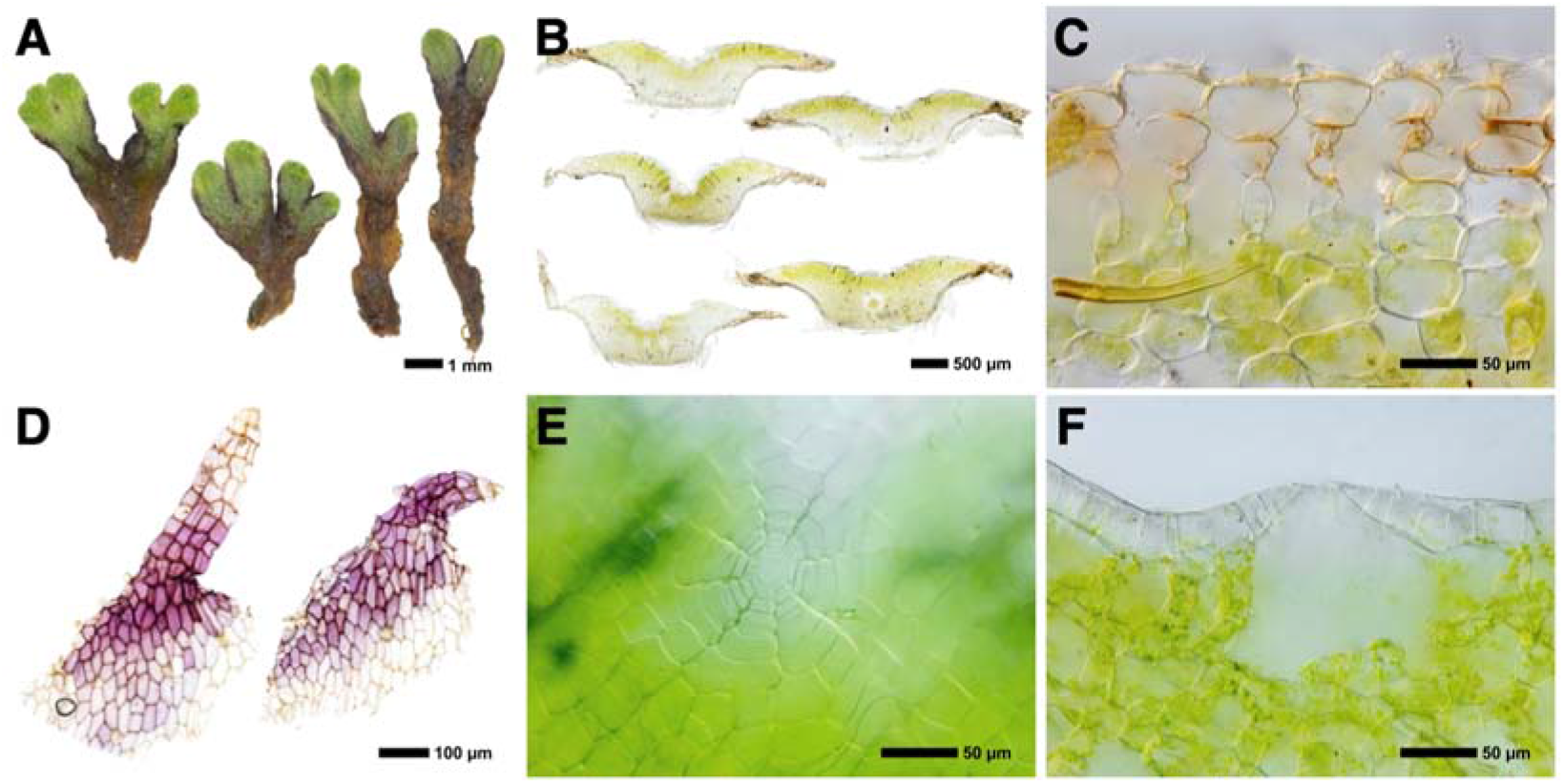
Exemplary morphological structures used for morphometric measurements listed in Supplemental Table S1. **(a)** Ventral (adaxial) side of thallus used for measuring width and length of the thallus. Image shows *Riccia gothica* at approx. 5x magnification. **(b)** Thallus slice in cross section used for measuring thallus width and height, and properties of the wing. Image shows *Riccia gougetiana* var. *armatissima* at approx. 50x magnification. **(c)** Epidermal and subepidermal cells in cross section. Image shows *Riccia bifurca* at approx. 400x. **(d)** Ventral scales that are violet pigmented and have slime cells. Image shows *Asterella gracilis* at approx. 200x. **(e)** Number of cells of air pore at anterior view. Image shows *Reboulia hemisphaerica* subsp. *hemisphaerica* at approx. 400x. **(f)** Air pore number of cells in cross section. Image shows *Reboulia hemisphaerica* subsp. *hemisphaerica* at approx. 400x.

**Table 1:**
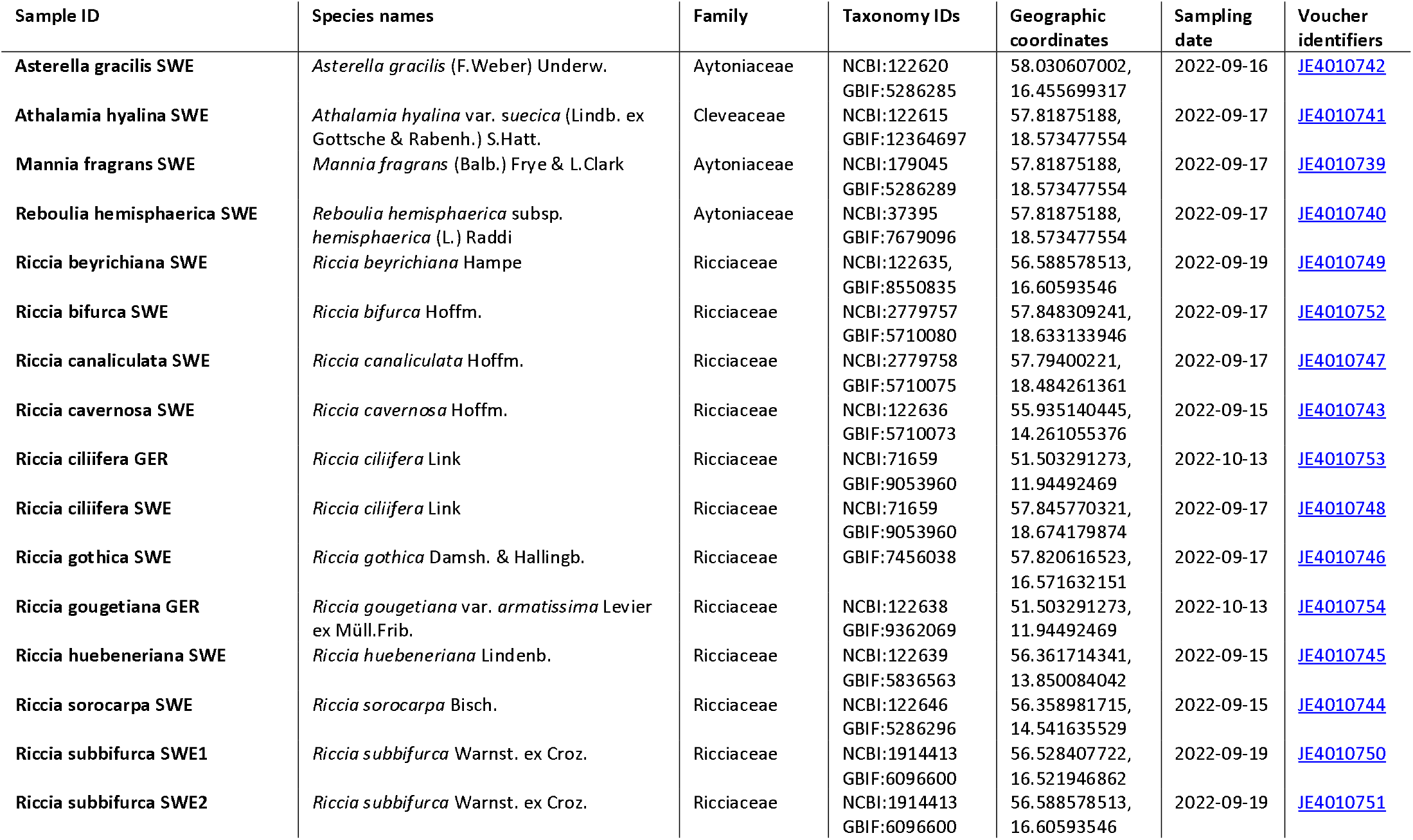
List of voucher specimens and fresh samples investigated in this study. The columns list the taxonomic species identifiers (*NCBI*, *GBIF*), the family the sample belongs to within the order of Marchantiales, geographic coordinates (WGS84 format, all coordinates have ± 10 m accuracy), sampling data and the voucher specimen identifiers of the Herbarium Haussknecht Jena (the first letters indicate the *Index Herbariorum* institution code^41^).

| Sample ID | Species names | Family | Taxonomy IDs | Geographic coordinates | Sampling date | Voucher identifiers |
| --- | --- | --- | --- | --- | --- | --- |
| Asterella gracilis SWE | *Asterella gracilis* (F.Weber) Underw. | Aytoniaceae | NCBI:122620<br>GBIF:5286285 | 58.030607002,<br>16.455699317 | 2022-09-16 | JE4010742 |
| Athalamia hyalina SWE | *Athalamia hyalina* var. *suecica* (Lindb. ex Gottsche & Rabenh.) S.Hatt. | Cleveaceae | NCBI:122615<br>GBIF:12364697 | 57.81875188,<br>18.573477554 | 2022-09-17 | JE4010741 |
| Mannia fragrans SWE | *Mannia fragrans* (Balb.) Frye & L.Clark | Aytoniaceae | NCBI:179045<br>GBIF:5286289 | 57.81875188,<br>18.573477554 | 2022-09-17 | JE4010739 |
| Reboulia hemisphaerica SWE | *Reboulia hemisphaerica* subsp. *hemisphaerica* (L.) Raddi | Aytoniaceae | NCBI:37395<br>GBIF:7679096 | 57.81875188,<br>18.573477554 | 2022-09-17 | JE4010740 |
| Riccia beyrichiana SWE | *Riccia beyrichiana* Hampe | Ricciaceae | NCBI:122635,<br>GBIF:8550835 | 56.588578513,<br>16.60593546 | 2022-09-19 | JE4010749 |
| Riccia bifurca SWE | *Riccia bifurca* Hoffm. | Ricciaceae | NCBI:2779757<br>GBIF:5710080 | 57.848309241,<br>18.633133946 | 2022-09-17 | JE4010752 |
| Riccia canaliculata SWE | *Riccia canaliculata* Hoffm. | Ricciaceae | NCBI:2779758<br>GBIF:5710075 | 57.79400221,<br>18.484261361 | 2022-09-17 | JE4010747 |
| Riccia cavernosa SWE | *Riccia cavernosa* Hoffm. | Ricciaceae | NCBI:122636<br>GBIF:5710073 | 55.935140445,<br>14.261055376 | 2022-09-15 | JE4010743 |
| Riccia ciliifera GER | *Riccia ciliifera* Link | Ricciaceae | NCBI:71659<br>GBIF:9053960 | 51.503291273,<br>11.94492469 | 2022-10-13 | JE4010753 |
| Riccia ciliifera SWE | *Riccia ciliifera* Link | Ricciaceae | NCBI:71659<br>GBIF:9053960 | 57.845770321,<br>18.674179874 | 2022-09-17 | JE4010748 |
| Riccia gothica SWE | *Riccia gothica* Damsh. & Hallingb. | Ricciaceae | GBIF:7456038 | 57.820616523,<br>16.571632151 | 2022-09-17 | JE4010746 |
| Riccia gougetiana GER | *Riccia gougetiana* var. *armatissima* Levier ex Müll.Frib. | Ricciaceae | NCBI:122638<br>GBIF:9362069 | 51.503291273,<br>11.94492469 | 2022-10-13 | JE4010754 |
| Riccia huebeneriana SWE | *Riccia huebeneriana* Lindenb. | Ricciaceae | NCBI:122639<br>GBIF:5836563 | 56.361714341,<br>13.850084042 | 2022-09-15 | JE4010745 |
| Riccia sorocarpa SWE | *Riccia sorocarpa* Bisch. | Ricciaceae | NCBI:122646<br>GBIF:5286296 | 56.358981715,<br>14.541635529 | 2022-09-15 | JE4010744 |
| Riccia subbifurca SWE1 | *Riccia subbifurca* Warnst. ex Croz. | Ricciaceae | NCBI:1914413<br>GBIF:6096600 | 56.528407722,<br>16.521946862 | 2022-09-19 | JE4010750 |
| Riccia subbifurca SWE2 | *Riccia subbifurca* Warnst. ex Croz. | Ricciaceae | NCBI:1914413<br>GBIF:6096600 | 56.588578513,<br>16.60593546 | 2022-09-19 | JE4010751 |

Even in large plant trait databases such as TRY, trait data of bryophytes are generally underrepresented mainly due to knowledge and methodological gaps^1,4,12^. The idea of molecular plant traits representing ecological functions or patterns via the presence/absence or abundance of small molecules has been around since the beginning of the century^13^. But only recently, they have been proposed to be integrated as conceptual entities into functional ecology^8^ mainly due to insufficient reference data and facilitated by recent computational breakthroughs in metabolomics such as *in silico* annotations^14,15^, *in silico* compound classification^16^ utilizing chemical ontologies^17,18^, computational molecular structure representations leading to quantitative structure-activity (QSARs) and -property relationships (QSPRs) and other chemical modelling approaches enabling the calculation of molecular descriptors^19–22^, computer-assisted community-guided chemical knowledge extraction^23^, open-source cheminformatics libraries^24^, or public libraries that compile and provide these various information using the FAIR principles^25,26^. However, data on molecular traits are still scarce, fragmented and ambiguous as controlled vocabularies and standardised frameworks are largely incomplete defining and naming the individual components of molecular traits relevant in ecology.

Here, we define Essential Molecular Variables (EMV). They are to functional ecology what Essential Biodiversity Variables (EBV) are to biodiversity^27^ – proxies that abstract from low-level primary observations and used as high-level indicators of functions (ecological observations). EMVs constitute only the most robust molecular traits and can be comprised of small molecules, compound classes (generalisation on molecule identity), or molecular descriptors (generalisation of physicochemical properties of molecules).

In order to improve the assessment of functional traits at different scales, here we present (1) reference data for functional ecology and biodiversity integrating three domains: (a) the assessment of phenotypic traits using bioimaging and morphometry, (b) the assessment of molecular traits using untargeted metabolomics (molecular computations performed on liquid chromatography high-resolution mass-spectrometry (UPLC/ESI-QTOF-MS) with data-dependent acquisition (DDA-MS) data), and (c) resolving phylogenetic traits by using DNA marker sequences. First, we utilize the bioimaging and metabolomics data and perform exemplary morphometric measurements and molecular computations demonstrating the principal extraction of phenotypic and molecular traits for use in the plant trait database TRY^12^. We further show how the data from the three different domains can be integrated, resolving evolutionary and ecological questions in integrative taxonomy and functional ecology. We (2) present the reference framework iESTIMATE to extract molecular traits and to estimate EMVs – those molecular traits that have high ecological relevance. We also propose formalized naming schemes. With this we want to encourage standardised and systematic cross-domain data integration, sharing, reuse and meta-synthesis in the fields of functional ecology, integrative biodiversity and related research disciplines.

## Results

### Computational framework

The iESTIMATE framework has been made available along with all re-usable pre-processed data as R package and can be installed in the following way:

devtools::install_github(“https://github.com/ipb-halle/iESTIMATE”)

Vignettes that explain how data is analyzed, dendrograms and other plots are made and traits are generated and formatted for TRY are available on GitHub.

https://github.com/ipb-halle/iESTIMATE/doc/

Figure 2 provides a schematic overview on the processing tasks of the iESTIMATE framework. The workflow starts with the experiment, where experimental sequence, phenotypic, or molecular data and metadata are being acquired for one or several biological species resulting in assembled and aligned sequences, a table with morphometric measurements, a table with molecular features and a table with molecular annotations (molecular structure, classification). The metadata is being used to acquire additional data from literature or external resources (i.e., based on species ID) and for the subsequent data analyses and estimations. Traits are extracted from the data tables and exported to TRY. Finally, data analyses (clustering and generation of dendrograms, calculation of diversity measures, variable selection) performed to reveal significant traits, and data integration (diagnostic plots, ordination, trait space calculation) to explore the quality of the generated trait data. Lastly, by a combination of various data analyses and variable selection, few essential molecular variables (EMVs) are distilled.

**Figure 2.**
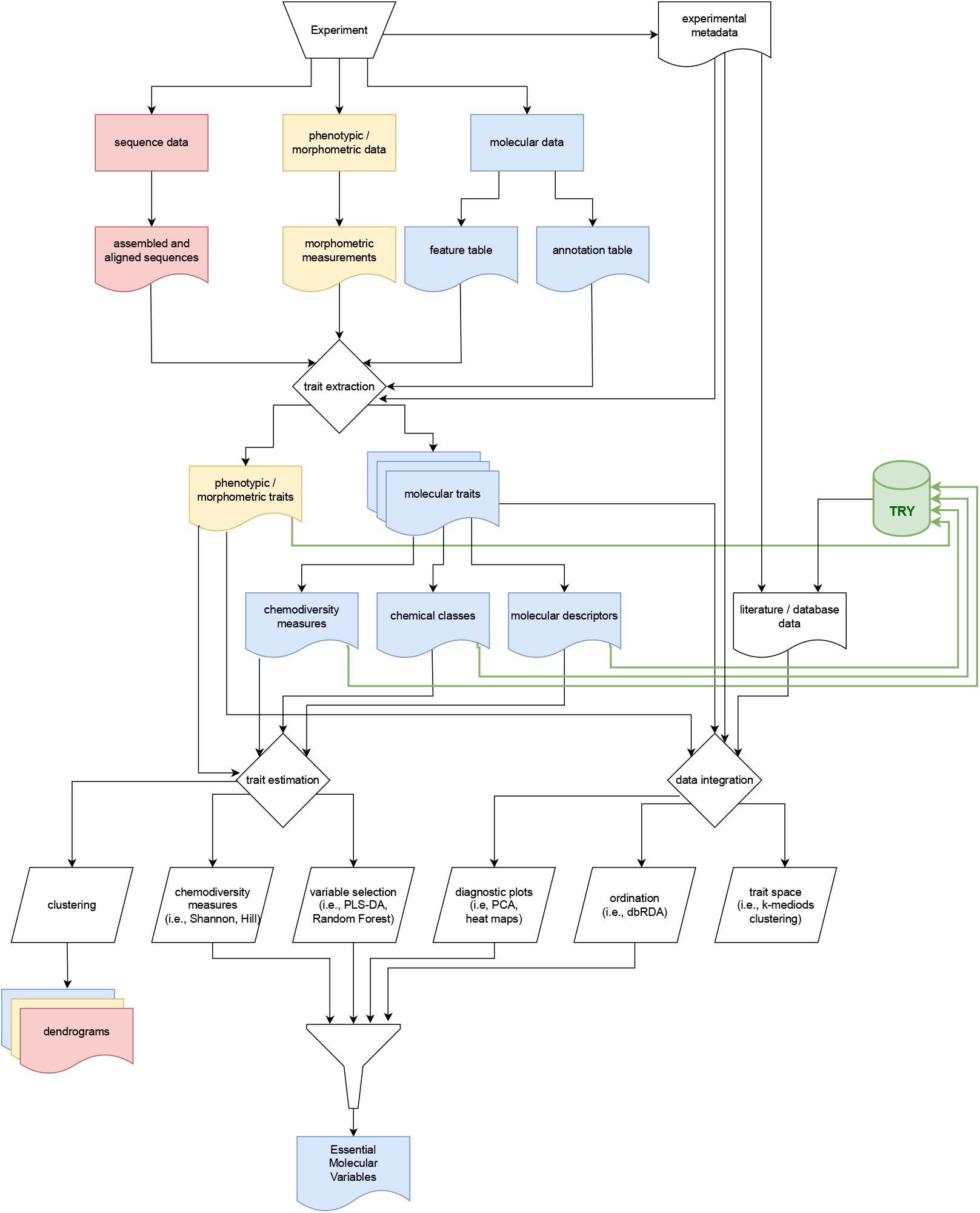
Flow chart for the iESTIMATE processing workflow. Based on input metadata, sequence-based data and phenotypic and molecular data in the form of matrices are used to perform trait extractions yielding phenotypic und molecular traits at different scales (chemodiversity measures, chemical classes and molecular descriptors). The traits can also be exported to TRY. Based on these traits, various analyses are performed, such as clustering revealing dendrograms, calculation and statistics of chemodiversity measures, unsupervised multivariate methods such as PCA, ordinations, k-medoids clustering for assessing trait spaces, and variable selection using PLS-DA and RF. Based on the data analyses, Essential Molecular Variables (EMVs) are finally estimated.

### Integration and synthesis of taxonomic data

To compare taxonomic data, we generated dendrograms separately for the sequencing data, molecular and phenotypic traits allowing for evolutionary inferences of the species reconstructed from the data at the respective level (Fig. 3). The similarity of the trees can be compared using the Robins-Foulds metric, i.e., by using the function RF.dist of the R package phangorn. The lower the value the more similar the trees. The similarity of the constructed trees can also be inspected visually (Fig. 3). Differences of the trees obtained at the molecular or phenotypic levels from the tree built from the sequences provide additional insights into species evolution and speciation that cannot be inferred from sequences alone, i.e., hybridization, trait multifunctionality, convergence, or ecological adaptation to environments or biotic interactions. In analogy to integrative taxonomy, integrating trait data at different levels can explain patterns and processes of diversity from multiple and complementary perspectives . Integrative trait data necessitate further investigation and can reveal functional processes at various ecological and evolutionary scales. Similar to dendrograms, the use of PCA plots were additionally proposed . However, both, dendrograms and PCA are only useful for explorative analysis. In order to make conclusions about the underlying eco-evolutionary mechanisms, we propose to first investigate the trait data at the distinct levels and to integrate the abstracted results subsequently. In summary, we find that molecular and morphometric trait data provide more information and resolution regarding species evolution than the *trn*LF marker and largely support species differentiation. However, the information can be confounded with location or community effects that need to be accounted for in ecological experiment designs, i.e., designing longitudinal studies containing more samples of the species in various locations. Exemplary code to perform integration of evolutionary data is available on GitHub (https://ipb-halle.github.io/iESTIMATE/doc/data_integration.html).

**Figure 3.**
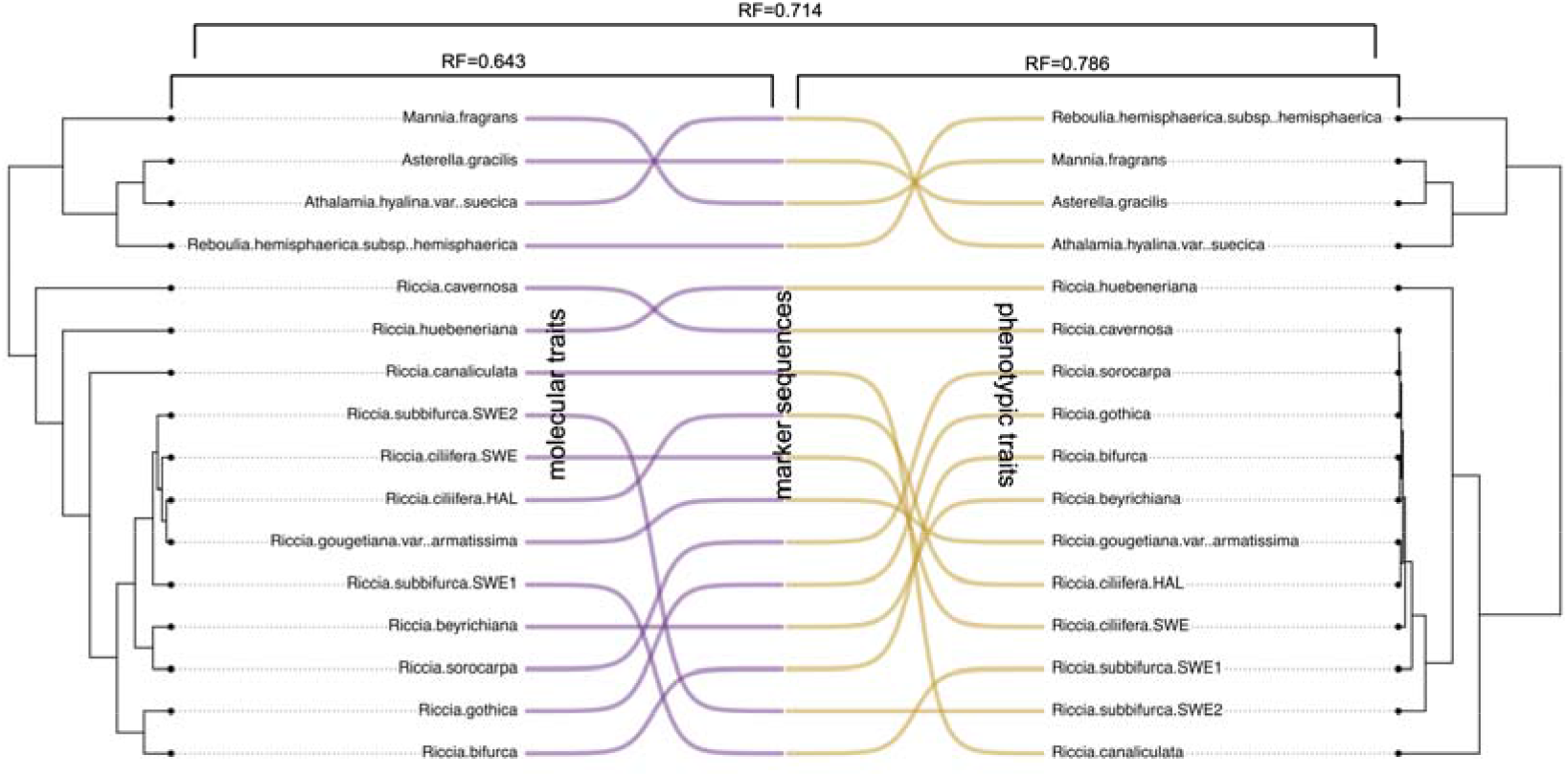
Comparison of dendrograms obtained from the selected molecular traits (at the level of compounds) (left, purple color), the trnLF sequences (center), and the measured phenotypic traits (right, dark yellow color). Species names and dendrogram resulting from markers are not shown in the center. Robinson-Foulds similarity of molecular traits and sequence trees: RF=0.643. Robinson-Foulds similarity of phenotypic traits and sequence trees: RF=0.786. Rebonson-Foulds similarity of molecular and phenotypic traits: RF=0.714. The lower the value the more similar the trees.

### Integration of functional traits

The Hutchinsonian multidimensional niche is an important concept in ecology that combines many ecological and evolutionary hypotheses^30^. It can be applied to functional ecology by estimating the hypervolumes that characterize the trait space occupied by a set of species. Evaluating the volume, distribution and overlap of species traits can explain how ecological and evolutionary mechanisms structure the functional responses of species^31,32^. This concept can be applied to phenotypic and molecular traits alike allowing for, both, integration of data and integration into functional ecology. To investigate the space of phenotypic and molecular traits, we combined k-medoids clustering^33^ of the respective trait matrices with subsequent 2D projection (Fig. 4a,b). In addition, we performed ordination using distance-based redundancy analysis (dbRDA) to correlate molecular with phenotypic traits and molecular with “classic” traits obtained from TRY^4^. Taken together, these analyses allow assessments of trait quality and gaining deeper insights into functional mechanisms^8^. Exemplary code to perform integration of functional ecology data is available on GitHub (https://ipb-halle.github.io/iESTIMATE/doc/data_integration.html).

**Figure 4.**
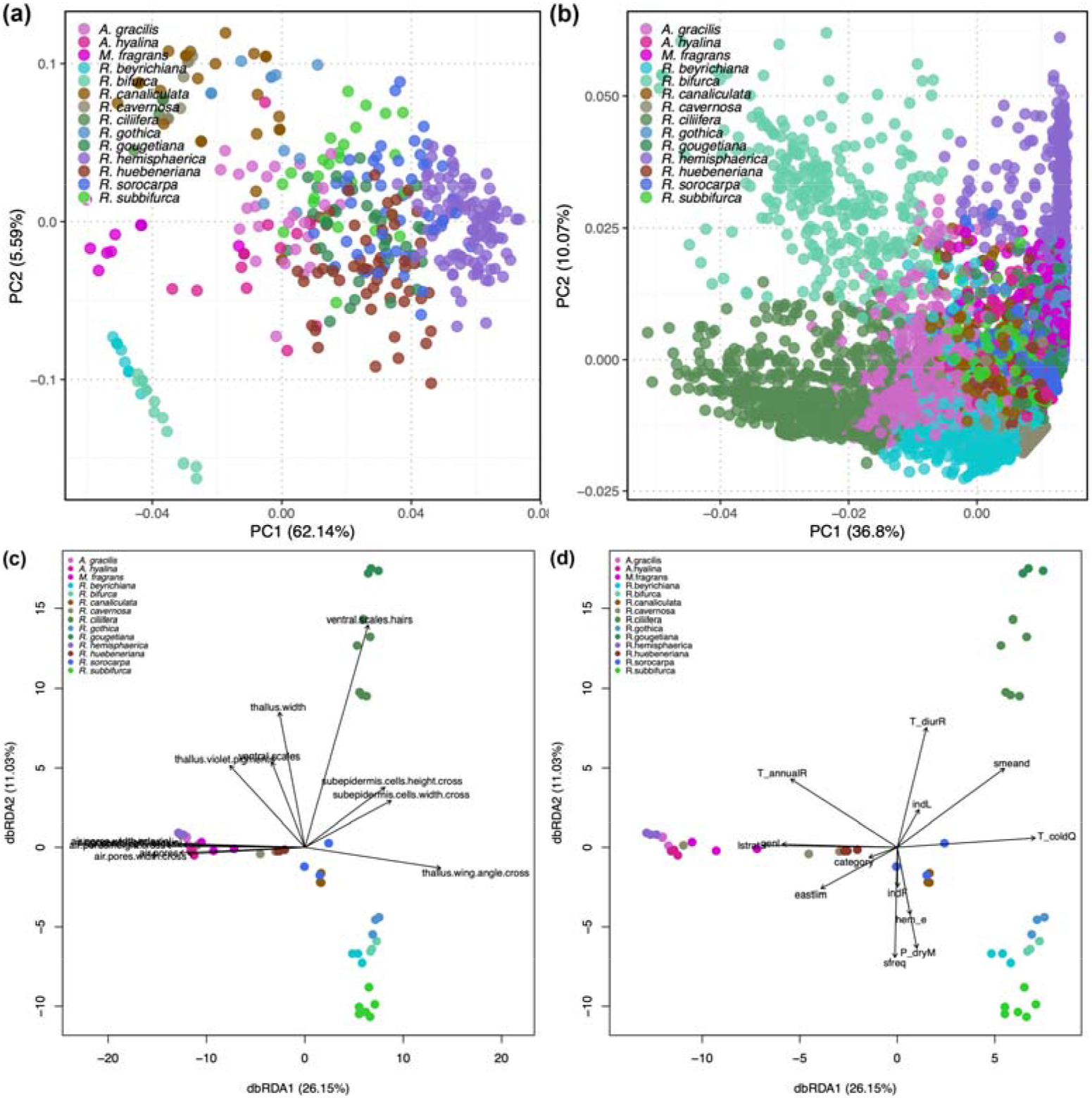
Hypervolumes shown as 2D projections for all combinations of (a) morphometric traits, and (b) molecular traits. To improve the morphometric model, hypervolumes for the morphometric traits were constructed by resampling measured traits 32 times. The more the loadings scatter the larger the theoretical trait space occupied by a species. Ordinations using distance-based redundancy analysis (dbRDA) of the (c) molecular traits with phenotypic traits, and (d) molecular traits with “classic” traits obtained from TRY. The length of the arrow represents the correlation of the respective trait with the molecular traits of species (shown by colored circles). The relative position of the species to the direction of the axis describes the relationship of the species with the respective trait.

### Estimation of Essential Molecular Variables (EMVs)

Inspecting our data resulted in 91 phenotypic traits (including variance, skewness and kurtosis calculations on the continuous morphometric measurements) and 46’462 individual molecular features. As molecular feature data is too complex to give meaningful results, data was filtered and only those features retained that correspond to molecules that had available spectral information. The matrix was further abstracted retaining only compound classes, molecular pathways and molecular descriptors. This resulted in molecular traits represented by 5’144 molecules, 455 classes, 140 subclasses, 16 superclasses^17^, 239 natural product classes, 7 natural product pathways^34^, and 241 molecular descriptors. In order to select for the EMVs underlying specific relationships, several different statistical or machine learning methods like PLS-DA, LDA, OPLS, Random Forest (RF), or SVM are available^35^. For demonstration, we use RF to select for the essential variables. Our framework also supports PLS-DA. Figure 5 shows exemplary heatmaps of EMVs at the levels of natural product class and pathway. Code to reproduce the results and common metrics like AUC, AUC-PR, R^2^, accuracy or balanced error rates to evaluate the model performance are available on GitHub (https://ipb-halle.github.io/iESTIMATE/doc/marchantiales.html).

**Figure 5.**
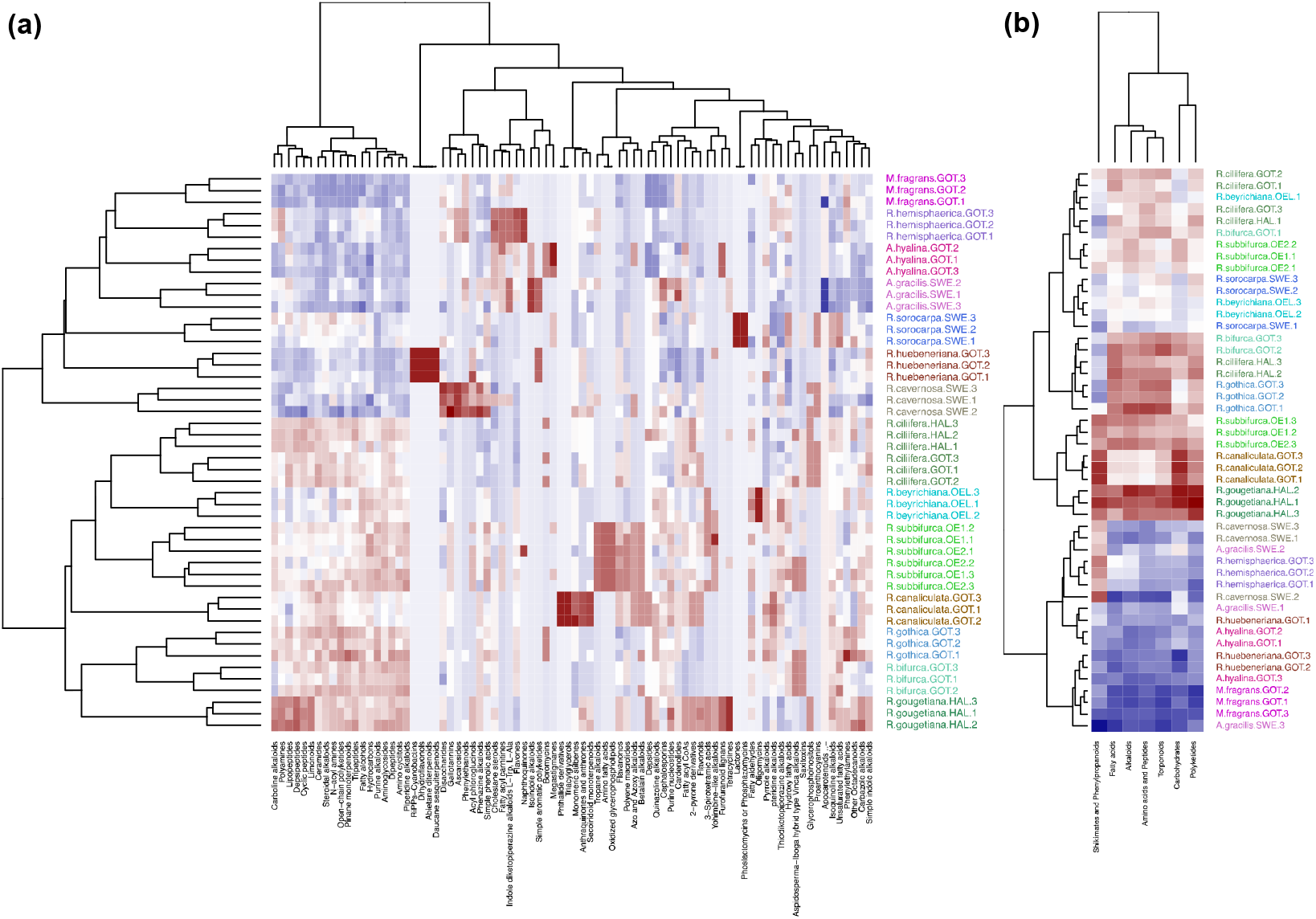
Heatmaps of EMVs at different abstraction levels. (a) EMVs at the level of natural product classes (82 selected variables, R^2^=0.970), (b) EMVs at the level of natural product pathways (7 selected variables, R^2^=0.233). Blue color means lower molecular count (underrepresentation) and red color higher count (overrepresentation).

## Discussion

We present the open-source computational framework iESTIMATE that reuses our reference data and subsequently integrates the data using established concepts in functional ecology and biodiversity (Fig. 2). iESTIMATE is reproducible, uses computationally lightweight methods and is well documented. We followed the FAIR principles^26^ and have designed our framework so that it can be used on other than our reference data. In principle, our modularized framework can also be formalized to computational workflows^36^. The described methods have the potential to increase the impact of shared integrative research data. In addition, our approach may be also useful in training programs as data integration is pivotal in many fields of biodiversity. With our iESTIMATE framework we want to encourage integrative data-driven research and data sharing in functional ecology and biodiversity. To this end, we have deposited raw and processed data in domain-specific repositories. We have made available all relevant processed reference data as data objects in the R package iESTIMATE and additionally individual tables in TSV format on GitHub.

Our framework iESTIMATE enables the extraction of phenotypic traits using morphometry and molecular traits at different levels of abstraction. As molecular data can be overwhelmingly complex, abstraction is needed in order to observe ecological relevant patterns and to make meaningful conclusions^37–39^. We support the arrangement of molecular traits into molecules (metabolic compounds and chemodiversity measures), compound classes (*in silico* classification utilizing the CHEMONT or NP-Classifier ontologies^17,18^), and molecular descriptors (generalizing on physicochemical properties of the molecules). We then support the application of variable selection methods such as RF and PLS-DA to select only for the Essential Molecular Variables (EMVs) that directly relate low-level primary observations to high-level ecological functions. They also facilitate data integration by providing an intermediate abstraction layer between primary observations and ecological functions^27^. Distilling the phenotypic and molecular data into EMVs and integrating them into a coherent ecological context enhances the explanatory power of the functional trait concept and offers a new set of tools for the discovery of mechanisms in functional ecology, biodiversity and related disciplines^2,3,8^.

## Methods

### Sample collection

Samples of were collected in the field and put immediately in sterile petri dishes at locations in Southern Sweden in September 2022 and in Germany in October 2022 (Table 1). Specimens were photographed on-site and bioimaging was performed on fresh material using macro- and microscopy in the lab. After an incubation period of five days at room temperatures, the plant material was isolated, washed under a light microscope to remove dirt and other residues, filled into Eppendorf tubes and shock-frozen (LC-MS analyses), or dried (marker sequencing). Remaining plant material was stored as voucher specimens in the Herbarium Haussknecht Jena (barcodes: JE04010739-JE04010754) (Table 1).

Validations of metadata, image properties and technical quality of fused composite images were conducted with the approach described in ^40^. Species identities were kindly validated by Tomas Hallingbäck and Galin Gospodinov.

### Image acquisition

For image acquisition, a Zeiss Axio Scope.A1 HAL 50, 6x HD/DIC, M27, 10x/23 microscope with an achromatic-aplanatic 0.9 H D Ph DIC condenser was used for microscopy utilizing the objectives EC Plan Neofluar 2.5x/0.075 M27 (a=8.8mm), Plan-Apochromat 5x/0.16 M27 (a=12.1mm), Plan-Apochromat 10x/0.45 M27 (a=2.1mm), Plan-Apochromat 20x/0.8 M27 (a=0.55mm), and Plan-Apochromat 40x/0.95 Korr M27 (a=0.25mm) using the EC PN and the Fluar 40x/1.30 III and PA 40x/0.95 III filters for DIC. The conversion filter CB3 and the interference filter wideband green were used to improve digital reproduction of colors. The color balance was adjusted in the camera and in software accordingly. For macroscopy and for preparing microscopy slides, a binocular stereo microscope Zeiss Stemi 2000c was used. For macroscopic images, the Venus Optics Laowa 25mm 2.5-5.0x ultra-macro for Canon RF was used. To acquire digital images, a full-frame, high-resolution camera (Canon EOS RP, 26 megapixel) was used and adapted to the photographic objectives or to the microscopes using binocular phototubes with sliding prism 30°/23 (Axio Scope.A1) and 100:0/0:100 reversed image (Stemi 2000c) using 60-T2 camera adapter for Canon EOS and a Canon EF-RF adapter. To construct images with extended depth-of-field, images were recorded at different focal planes and by attaching the camera to a Cognisys StackShot macro rail fixed on a Novoflex macro stand, and for microscopy by adapting a Cognisys StackShot motor to the fine adjustment of the microscope using two cogged wheels, one small wheel (1 cm diameter) adapted on the motor and one large wheel (8.5 cm diameter) on the fine adjustment of the microscope. The two cogged wheels were coupled with a toothed belt to obtain fine step increments of the stepping motor for high magnifications. A Cognisys StackShot controller was used to control the amount and distance of the stepping motor with the following controller settings: Dist/Rev: 3200 stp, Backlash: 0 steps, # pics: 1, Tsettle: 100.0 ms, Toff: 450.0 ms, Auto Return: yes, Speed: 3000 st/sec, Tlapse: off, Tpulse: 800.0 ms, Tramp: 100 ms, Units: steps, Torque: 6, Hi Precision: Off, LCD Backlight: 10, Mode: Auto-Step using between 25 steps (magnification 1x) and 50 steps (magnification 25x) and 100 steps (magnification 400x) (number of steps depending on aperture settings and effective magnification).

### Image processing

Raw images were recorded in CR3-format and pre-processed with Adobe Lightroom Classic 2022 where non-destructive image processing such as corrections of the field curvature, removal of chromatic aberration, color balance, increase of contrast and brightness were performed^42^. Images were then exported to TIFF-format and any image processing steps were recorded in individual Adobe XMP-files. Multi-focus image fusion was performed on the individual images in the z-stacks using the software Helicon Focus 8.2.9 and by choosing the algorithms depth map and pyramid with different settings of radius (4, 8, 16, 24) and smoothing (2, 4, 8). The best composite image was chosen manually and retained. When composite images contained specimen that were larger than the frame, several images were stitched together using the Photomerge-Reposition function in the software Adobe Photoshop 2023. Images were manually segmented and interfering background removed using the object selection tools. In order to determine the scale, a stage micrometer was photographed separately with any of the objectives and microscope combinations. The scale was calculated per pixel for each combination and scale bars were put post-hoc onto the segmented images.

### Extraction of phenotypic traits: Morphometric measurements

Measurements of morphometric characters were performed manually with Fiji^43^. After setting up the scales for the individual images in the Image-Properties menu, the following morphometric characters were measured: thallus width [µm], thallus length [µm], thallus with violet pigments [0/1], ventral scales [0/1], ventral scales with slime cells [0/1], ventral scales with violet pigments [0/1], ventral scales with hairs [0/1], air pores [0/1], width of ring cells of air pores in adaxial view [µm], height of ring cells of air pores in cross section [µm], number of ring cells of air pores in cross section [#], width of ring cells of air pores in cross section [µm], height of ring cells of air pores in cross section [µm], width of epidermis cells in cross section [µm], height of epidermis cells in cross section [µm], width of subepidermal cells in cross section [µm], height of subepidermal cells in cross section [µm], width of thallus in cross section [µm], height of thallus in cross section [µm], height of thallus wing in cross section [µm], angle of thallus wing in cross section [°], width of thallus wing in cross section [µm], area of thallus in cross section [µm^2^] (Figure 1, Supplemental Table S1). Lengths and widths were obtained using the Measure function from the Analyze menu and the results saved in CSV files. To automate the measurement of area, a pixel classification model was trained using LabKit^44^ by selecting representative foreground and background areas. The imaging data was segmented subsequently with StarDist^45^ using the above classifier. StarDist automatically classified objects in the images into foreground and background and areas classified as foreground were then measured using the Measure function from the Analyze menu and results were saved. CSV files with all individual morphometric measurements of all specimens were joined into one single table and used for subsequent data analyses.

In order to represent the high plasticity and variability of morphological characters of liverworts, variance, skewness and kurtosis of any of the measured continuous morphometric characteristics (Supplemental Table S1) were calculated additionally according to ^46,47^.

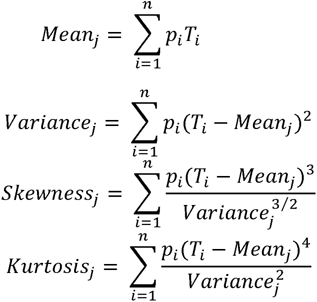

For each character, *p*_i_ represents the measurements for species *j*, *T*_i_ the mean value of character measurements of all species, *n* is the total number of measurements for the character. The variance and skewness represent the dispersion range and asymmetry of the character distribution, respectively. The kurtosis indicates the relative peak of the character distribution and the heaviness of its tails. Low kurtosis reflects high evenness in abundance of character values which implies similarity of species.

### Image data management

Metadata including species name, taxonomic rank information (NCBI-Taxon and GBIF taxonomy identifiers), voucher specimen id, image acquisition date, an object description including the name of the captured morphological character(s), the used objective, microscope, and magnification were associated with any raw image based on unique respective file names. Individual file names, name within an image focus stack and name within an image stitching stack were recorded additionally to facilitate subsequent automated image processing in workflows. Python scripts were created to automate image fusion and image stitching tasks. Raw camera and pre-processed imaging data in CR3 and TIFF format, respectively, were deposited to the BioImage Archive (BioStudies) using the command line IBM Aspera software tool ascp version 3.8.1.161274. The raw bioimaging data is available under the BioStudies identifier S-BIAD824 (https://www.ebi.ac.uk/biostudies/studies/S-BIAD824). Processed images were converted to the Bio-Formats OME-TIFF format by creating intermediate ZARR-pyramid tiles using the bioformats2raw converter version 0.7.0 and then using the raw2ometiff version 0.5.0 software tool to create the final pyramid images. Processed images and the metadata were first aggregated in a TSV table and then deposited to the Image Data Resource (https://idr.openmicroscopy.org/search/?query=Name:157) under accession number idr0157 using the software Globus Connect Personal 3.1.6.

### DNA marker sequencing

DNA marker sequences of the *trn*L-*trn*F (*trn*LF) plastid region were obtained by the German Barcode of Life using liverwort-specific primers described in ^48^. Sequences were analyzed using an AB 3730 DNA analyzer instrument and PeakTrace was used as basecaller and trace processor. Multiple sequences were aligned using the MUSCLE 3.8 REST web service Perl client^49^. The resulting FASTA file was used for further phylogenetic analyses.

### Sequencing data management

Sequencing data was deposited to the European Nucleotide Archive (ENA)^50^ and is available under the study identifier ERP155252 (accession PRJEB70317) (https://www.ebi.ac.uk/ena/browser/view/PRJEB70317). Raw reads of *trn*LF sequences of the 16 samples are available under the sample identifiers SAMEA114863468-SAMEA114863483.

### Construction of dendrograms

The phylogenetic reconstructions for the plastid region of *trn*LF were conducted with RAxML-NG 1.2.0^51^ using the model GTR with discrete GAMMA (GTR+G). Alpha of phylogenetic trees was estimated using Maximum Likelihood (100) and clade support were estimated using bootstrapping (1000). Additional nucleotide sequences of the investigated species were obtained from Genbank^52^, if available, using the R package rentrez and included in the construction of the phylogenetic tree. Values of phenotypic traits (morphometric measurements) and molecular traits were discretized into 8 states using the gap weighting algorithm^53,54^. Phylogenetic reconstructions of the resulting data were accomplished using RAxML-NG, but with using a multistate model (MULTI8_GTR). Dendrograms were constructed in R using the plotBS function of the R package phangorn. Co-phylogenetic plots were constructed using the cophylo function from the phytools package. Exemplary code to produce trees is available on GitHub (https://ipb-halle.github.io/iESTIMATE/doc/data_integration.html).

### Validation of marker sequence data

For many liverworts (division Marchantiophyta) it is challenging to obtain DNA marker sequences like nuclear ITS mainly due to high frequencies of repetitive elements and a high abundance of polyphenols, flavonoids, fatty acids and other specialised metabolites that coprecipitate to DNA fragments with common DNA and RNA extraction procedures^55^. In order to validate species identity, chloroplast *trn*LF marker sequences were obtained herein and compared with those available in Genbank. To this end, a dendrogram was constructed containing 16 sequences of the investigated samples and 33 sequences obtained from Genbank (Figure 6). For the species *Riccia bifurca* , *R. canaliculata* , *R. ciliifera*, and *R. gothica* no sequences were available in Genbank. The species *R. ciliifera* (GER) and *R. gougetiana* (GER), as well as *R. beyrichiana* (SWE-Öland) and *R. bifurca* (SWE-Gotland) could not be differentiated with the obtained *trn*LF sequences due to the limited resolution of the marker. Our results indicate that the deposited sequence of KX468755.1 *R. subbifurca* could be *R. gothica* instead, or the species identities of the herein investigated *R. subbifurca* and *R. gothica* were interchanged despite the fact that morphological investigation of spores indicated otherwise. All other samples cluster at expected positions in the tree. In summary, the plastid marker *trn*LF does not provide sufficient resolution to unequivocally differentiate the investigated species. Seeing the difficulties of obtaining more informative marker sequences, alternative methods such as the use of morphological characters or chemometric markers using metabolomics are recommended (see below). The comparative sequence analysis is available on GitHub (https://ipb-halle.github.io/iESTIMATE/doc/data_integration.html).

**Figure 6.**
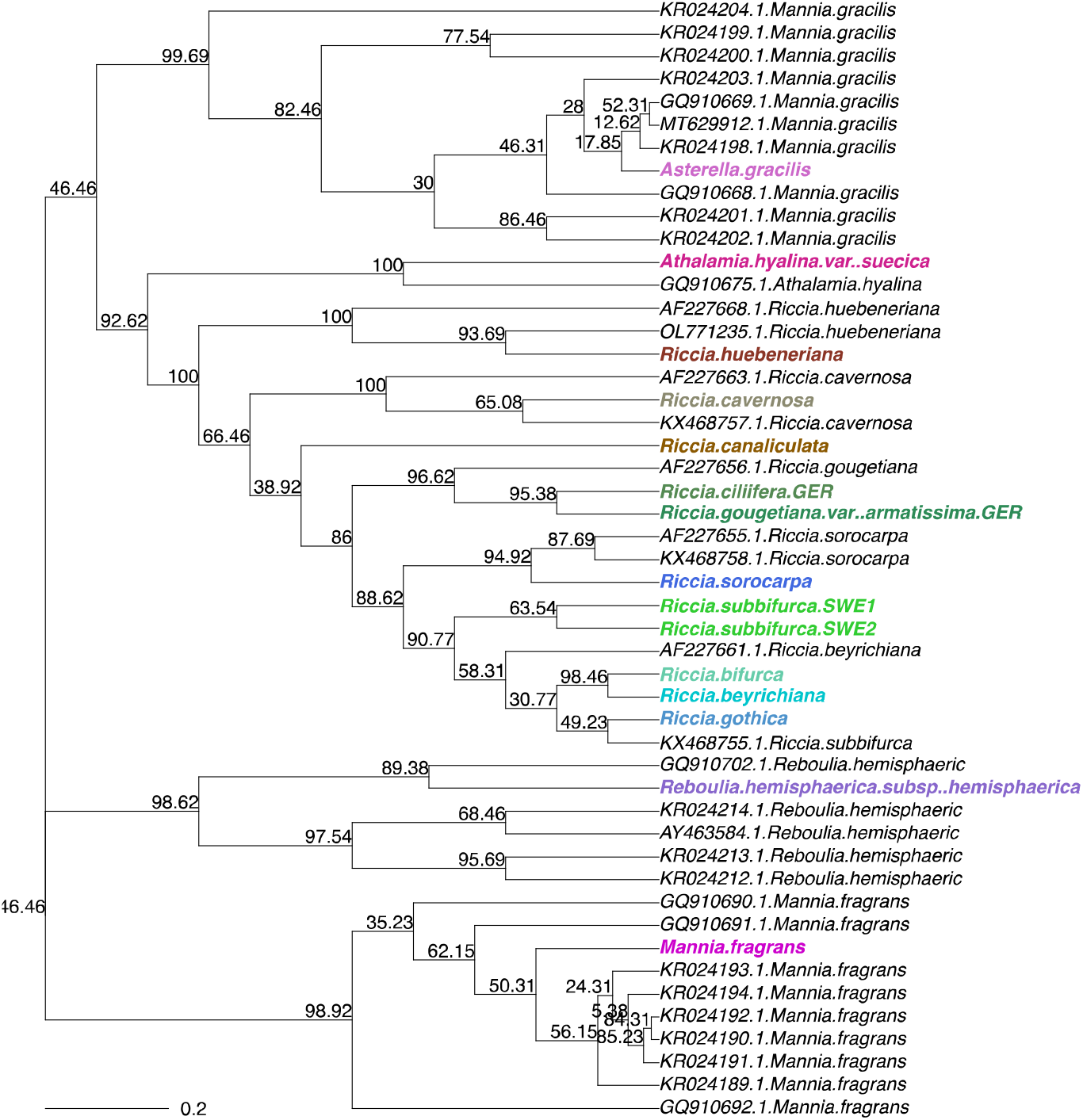
Phylogenetic tree of the *trn*LF marker sequences of the investigated species samples and those available from Genbank. Investigated species in bold and color.

### Metabolite extractions

We followed extraction procedures for LC-MS originally developed for vascular plants by ^56^ and adapted for the use with bryophytes and the different analytical setup used herein by ^57^. This data-dependent acquisition method has been shown to provide robust results and high-quality spectra for thallose liverworts^58^. In detail, frozen plants were homogenized for 5 minutes at 30 Hz in Teflon chambers equipped with two stainless steel balls. The chambers were rinsed with 100 µL of cold 80:20 MeOH:H2O supplemented with 2 µM 4-Methylumbelliferon, 5µM Kinetin (Sigma), 5µM Biochanin A (Sigma), and 5 µM N-(3-Indolylacetyl)-L-alanine (Sigma) and the slurry was shaken at 1000 rpm for 15 minutes at room temperature. The mixture was centrifuged twice at 13000 rpm for 15 minutes and the supernatant was transferred to autosampler vials.

### Untargeted LC-MS-MS Analysis

Samples were separated with an Agilent 1290 Infinity HPLC (Agilent, Waldbronn, Germany) equipped with a Nucleodur X18 Gravity-SB column (1.8µm 100x2 Macherey Nagel, Dueren, Germany) coupled to a Bruker TIMS-TOF-MS (timsTOF Pro, Bruker, Bremen, Germany). Separations were performed at 35°C with the following binary gradient of 0.1% aqueous formic acid (solvent A) and acetonitrile (solvent B): isocratic 98% A for 1 minute; a linear gradient from 98% to 5% A from 1 to 14 minutes; isocratic 5% A from 14 to 17.5 minutes; isocratic 98% A from 18 to 20 minutes. The flow rate was maintained at 0.5mL/min, the injection chamber was maintained at 4°C. Injections were performed separately for positive and negative ionization mode. Data-dependent acquisition (DDA-MS) mode was used with the instrument settings described in ^57^.

### Raw data and MS1 data processing

Raw data converted into mzML format using msconvert^59^ as well as derived data (SIRIUS project folders, RData) were deposited in MetaboLights under the study identifier MTBLS2239^60^. Metadata were recorded in compliance with the minimum information guidelines for Metabolomics studies^61^.

Data processing was performed in the statistical software environment R version 4.3.1 using the iESTIMATE framework (https://github.com/ipb-halle/iESTIMATE). Chromatographic peak detection was performed using the R package XCMS version 3.22.0^62^. The following settings were used for the negative ion mode: CentWaveParam, ppm = 25, mzCenterFun = “mean”, peakwidth = c(7.2, 36), prefilter = c(2, 50), mzdiff = 0.0012, snthresh = 7, noise = 0, integrate = 1, firstBase-lineCheck = TRUE, verboseColumns = FALSE, fitgauss = FALSE, roiList = list(), roiScales = numeric()). For the positive ion mode different settings were used for: peakwidth = c(9.4, 32), prefilter = c(6, 51), mzdiff = -0.0043, snthresh = 2. Grouping of chromatographic peaks was performed before and after retention time correction with the following settings in both ion modes: PeakDensityParam, minFraction = 0.7, bw = 0.25, minSamples = 1, binSize = 0.5. Retention time correction was performed with the following settings: PeakGroupsParam, minFraction = 0.7, smooth = “loess”, span = 0.2, family = “gaussian”. Only metabolite features with retention times less than 1020 s were considered for further analysis.

The MS1-level peak tables were created separately for positive and negative ion modes with the settings featureValues, method = “medret”, value = “into”. The peak tables were log2-transformed, and missing values were imputed with zeros. Histograms and PCA diagnostic plots were generated to additionally evaluate the distribution of the data. MS2-level fragment spectra (MS-MS spectra) that were acquired by the Data-Dependent Acquisition mode (DDA-MS) were extracted from the profiles using the chromPeakSpectra, msLevel = 2 L, return.type = “Spectra” settings of XCMS. Spectra obtained from the same precursor ion were combined using the combineSpectra function from the R package Spectra using the following settings: FUN = combinePeaks, ppm = 25, peaks = “union”, minProp = 0.8, intensityFun = median, mzFun = median, backend = MsBackendDataFrame. These steps were performed separately for positive and negative ion modes. The MS1-level peak tables were then filtered to include only peaks for which the DDA-MS had acquired MS-MS fragment spectra. The spectra were saved in MSP and MGF files for further data processing. The peak tables and associated spectral and annotation data for positive and negative modes have been made separately available in MetaboLights.

As standard variance and median values were below 5% deviations, the filtered MS1-level peak tables containing log-transformed abundances of peaks in positive and negative ion modes were joined and used for further statistical analyses. Presence/absence peak tables were also generated to contain whether a metabolite feature was detected in the profiles. Features with abundances less than 10^−8^ % of the median abundance were considered absent. Peak detection R scripts are available for positive and negative modes on GitHub (https://github.com/ipb-halle/iESTIMATE/tree/main/use-cases/marchantiales).

### Annotation of MS-MS spectra

Annotation of MS-MS fragment spectra was carried out using the software SIRIUS version 5.7^14^. The settings described in ^58^ were used for both ionizations. Annotation was accomplished automatically by selecting the highest-ranking candidate for each spectrum. If the software could provide a COSMIC score^63^, the candidate with the highest-ranking COSMIC score was selected. The corresponding SMILES and the compound classification provided by the CANOPUS^16^ were extracted and stored for each spectrum. The classification provided by CANOPUS for each MS-MS fragment spectrum was aggregated and stored in a separate classification table. Compound classes were analyzed at the CHEMONT level of subclasses and superclasses. The classes were aggregated and counted for each spectrum found in a sample and multiplied by the (log2-transformed) peak abundances of the corresponding MS1 precursors in the MS1-level peak table described above. The same procedure was accomplished for classes obtained from the Natural Product Classifier^18^. Molecular descriptors were calculated for the SMILES provided by SIRIUS using rCDK^64^ resulting in a data matrix with SMILES in rows and descriptors in columns. A data table was constructed corresponding to the feature table by performing a matrix operation of both tables. This data table was used for performing the data analyses (see below). R scripts that annotate spectra are available for positive and negative modes on GitHub (https://github.com/ipb-halle/iESTIMATE/tree/main/use-cases/marchantiales).

### Extraction of chemodiversity traits

To assess the overall chemical diversity, the molecular richness was first determined representing the number of molecular features, molecules, classes, and descriptors found in a sample, respectively. Second, the number of unique variables was determined that represents those variables that are present in one species but not the others. As a third diversity measure, the Shannon diversity index H’ was determined according to ^10^. Finally, the Pielou’s evenness J that describes the homogeneity of the distribution of the intensity or abundance of compounds present in a species was determined according to ^10^. To assess significant differences among the groups, ANOVA with post-hoc Tukey honestly significant difference (HSD) test was calculated, and the R packages vegan, multcomp, and multtest were used. To get an overview of the chemical diversity of classes at different levels and their diversity among or across species, sunburst plots were constructed^10^. They were implemented as a custom function comprised as stacked barplots from the inside out, starting with organic compounds in the center^65^. The classes further to the outside represented the more specialized classes. The classes were arranged at different levels based on the CHEMONT or NP-Classifier ontologies^17,18^.

### Extraction of molecular traits

For explorative unsupervised multivariate analyses, principal components analysis (PCA) was performed using the prcomp function in R. In order to assess the influence of different study factors, variation partitioning was performed using the function varpart in the package vegan.

Variables were selected at the levels of MS1 features (“feature list”), MS1 features constrained to the availability of MS2 spectra (“molecule list”), at the subclass, class and superclass levels (“subclass list”, “class list”, “superclass list”) of the chemical ontology^17^, at the pathway and class level (“nppathway list”, “npclass list”) of the natural product ontology^18^, and at the level of molecular descriptors (“descriptor list”). Variable selection was accomplished with Random Forest (RF) using the caret package. A prediction model was trained using the train function from the caret package, and variable importance values were extracted from the model using the varImp function. Variables were selected (hence, were considered significant) when their quantile threshold was above 0.95. In order to visualize significant relationships of the selected variables at the different levels, heatmaps were generated from the selected variables using the gplots R package. To evaluate the performance of the fitted models, 10-fold cross-validation was performed (package mltest), and the Receiver Operating Characteristic (ROC) and PR (Precision and Recall) curves using the functions plot.roc and ci.se from the pROC package and the function pr.curve from the PRROC package were additionally constructed^66,67^. The R-squared of the fitted vs. the entire model and the area under curve (AUC) were calculated from the ROC, and the area under precision recall curve (AUC-PR) was determined from the PR curve.

### Metabolomics data management

Raw metabolite profiles and the annotated feature tables were deposited in the MetaboLights repository (study identifier MTBLS2239). Code to reproduce the results and generating the MAF for MetaboLights is available on GitHub (https://ipb-halle.github.io/iESTIMATE/doc/marchantiales.html).

### Integration of bioimaging and metabolomics data

To assess trait spaces at phenotypic and molecular scales, we used the pam function from the R package cluster and the autoplot function from the ggplot2 package. To get a more complete representation of the phenotypic trait space in comparison to the molecular trait space, trait measurements were resampled 32 times.

To investigate relationships of the phenotypic traits (morphometric measurements) and the molecular traits (small metabolic molecules), distance-based ReDundancy Analyses (dbRDA) were performed using the dbrda function of the package vegan. A Euclidean distance measure was used for the ordination. The dbRDA-model with the largest explained variance was chosen using forward variable selection and the ordistep function. The goodness of fit statistic (squared correlation coefficient) was determined for the remaining variables by applying the envfit function on the dbRDA ordination model. Code to reproduce the results is available on GitHub (https://ipb-halle.github.io/iESTIMATE/doc/data_integration.html).

### Estimation of Essential Molecular Variables (EMVs)

To select for the essential molecular variables (EMVs), a Random Forest (RF) supervised learning and classification method was carried out using the caret and randomForest R packages. A prediction model was trained using the train function from the caret package, and variable importance values were extracted from the model using the varImp function. Selected variables were considered significant when their variable importance was > 0.95. To evaluate the performance of the RF models, 10-fold cross-validation with 5 repeats was performed and the Receiver Operating Characteristic (ROC) and PR (Precision and Recall) were evaluated using the functions plot.roc and ci.se from the pROC package and the function pr.curve from the PRROC package^66–69^. The R^2^ of the fitted vs. the entire model and the area under curve (AUC) were calculated from the ROC, and the area under precision recall curve (AUC-PR) was determined from the PR curve. In order to visualize significant relationships of the selected variables at the different levels, heatmaps were generated using the gplots R package and interactive heatmaps using the heatmaply R package. Code to reproduce the results and the model metrics are available on GitHub (https://ipb-halle.github.io/iESTIMATE/doc/marchantiales.html).

### Extraction of morphometric traits for TRY

The morphometric traits listed in Supplemental Table S1 were measured in Fiji using the approach described above. Measurements were joined into one single table and imported into R. Variance, skewness and kurtosis were calculated for the measured variables with continuous (numeric) values. The resulting data table was exported with the traits listed in Table S1 to TRY.

### Estimation of molecular traits for TRY

Molecular traits in TRY were first represented by simple chemodiversity measures on the MS1 feature table characterizing the chemical diversity of species samples (Supplemental Table S2a).

Molecular traits in TRY are then represented by compound classes. Here, we preferred NP-Classifier over CHEMONT as it provides a more biologically meaningful interpretation^18^. The NP-Classifier is composed of molecular pathways (7 entities), superclasses (70 entities) and classes (672 entities)^34^. In TRY, the high-level information is based on molecular pathways and the low-level information on the molecular class (Supplemental Table S2b). In the investigated liverwort species, we annotated a total of 206 classes. Only approx. half of the theoretical classes defined in NP-Classifier can be found in plants.

Lastly, molecular traits were also represented by 241 molecular descriptors – physicochemical properties calculated on any annotated molecular structure of the individual species. Supplemental Table S2c lists the high- and low-level molecular descriptors calculated using rCDK. The high-level information is a summarised, abstracted representation while the low-level information consists of the CDK nomenclature.

### Export of estimated traits to TRY

Exemplary code to format and export traits of different kinds to TRY is available on GitHub (https://ipb-halle.github.io/iESTIMATE/doc/trait_export.html).

### Usage Notes

Raw and processed data, and meta-data are available under the terms of the Creative Commons licenses. Open-Source software scripts and code are available under the terms of the BSD 3-Clause license.

### Data Records

Two separate data records were created in order to enable rapid use of the data in machine learning and biodiversity approaches.

1. The camera raw images (Canon CR3-format), the pre-processed images (16-bit TIFF-format) and the contextual metadata were deposited to BioStudies under the identifier S-BIAD824 (https://www.ebi.ac.uk/biostudies/studies/S-BIAD824). The data record consists of a total of 223’989 individual raw image files partitioned into 48 species. The entire data record has a total size of approx. 12 TB. The pre-processed and processed images along with metadata were deposited to the Image Data Resource (IDR) repository under the identifier idr0157 (https://idr.openmicroscopy.org/search/?query=Name:157). The data record consists of a total of 4233 pre-processed and 905 fully processed imaged files. The data record has a total size of approx. 14 TB. Fully segmented images arranged by voucher specimens are available in Zenodo (https://doi.org/10.5281/zenodo.10683968).
2. Raw metabolite profiles (zipped vendor data and in mzML format) have been made available in MetaboLights under the study identifier MTBLS2239 (https://www.ebi.ac.uk/metabolights/MTBLS2239/). The dataset includes 48 metabolite profiles in positive and negative modes, QC and blank profiles, metabolite feature tables (MAF) and metadata.
3. Sequencing data were deposited to the European Nucleotide Archive (ENA) and are available under the study identifier ERP155252 (accession PRJEB70317) (https://www.ebi.ac.uk/ena/browser/view/PRJEB70317). Raw reads are available under the sample identifiers SAMEA114863468-SAMEA114863483.
4. Extracted ecological traits were submitted to the plant trait database TRY (https://www.try-db.org).
5. Metadata to voucher specimens is available at JACQ with the following identifiers: JE4010742, JE4010741, JE4010739, JE4010740, JE4010749, JE4010752, JE4010747, JE4010743, JE4010753, JE4010748, JE4010746, JE4010754, JE4010745, JE4010744, JE4010750, JE4010751.

## Code availability

The iESTIMATE framework containing functions, code, scripts and reference data is available as Open Source on GitHub (https://github.com/ipb-halle/iestimate). Scripts used in this study to process the imaging data are available on GitHub ( https://github.com/korseby/create_image_stacks, https://github.com/korseby/scale_bar, https://github.com/korseby/bioimage_submission). Python scripts were tested under Python 3.9. R scripts were tested under R 4.3.1. Shell scripts were tested using Bourne Again Shell (bash) 5.2.21.

## Acknowledgements

KP acknowledges the support of iDiv (funded by the German Research Foundation, DFG-FZT 118, 202548816). We would like to thank Tomas Hallingbäck and Galin Gospodinov for help with identification of species, and Nils Cronberg for help with coordinating the sampling campaign in Sweden. We would also like to thank the German Barcode of Life (GBOL) initiative for sequencing of DNA markers.

## Author contributions

KP conceptualised and performed the entire study. JZ conducted metabolomics extractions and measurements. SN acquired funding for the study and reviewed and edited the final draft.

## Competing interests

No competing interests.

Bibliographic information for any works cited in the above sections, using the standard *Nature* referencing style.

## Supplemental Material

### Technical validation of metabolomics data

The 48 samples were run in randomized injection order in one instrument run to avoid batch effects. Diagnostic plots of XCMS were employed to validate instrument performance and to detect shifts between the instrument runs. First, raw chromatograms were visually inspected for any shifts in intensity (Fig. S1, first row). Next, retention time (RT) correction made by XCMS on the samples was assessed (Fig. S1, second row) followed by mass-to-charge deviation (m/z) (in ppm) (Fig. S1, third row) and RT deviation (in seconds) (Fig. S1, fourth row). Lastly, total ion current (TIC) was determined and compared for the samples (Fig. S1, last row). In summary, we found maximum retention time deviations within 2 seconds – which is well within limits of the analytical setup used. The determined maximum mass-to-charge deviations of 4 ppm (negative ion mode) and 6 ppm (positive mode) were within instrument specification as well. For the majority of samples, the deviations were even less than 1 ppm (Fig. S1, third row). Deviations in mass-to-charge and retention times were corrected by XCMS. We conclude that there are no significant shifts in the instrument run and that the corrections made by XCMS are validated for the parameters used for peak detection.

In addition, the intensities of three internal lab standards that were spiked into all samples were also investigated (Fig. S2). We found that the variation for the internal lab standard were within the typical range of 10-15%^52^.

### Estimation of morphometric traits

**Figure S1.**
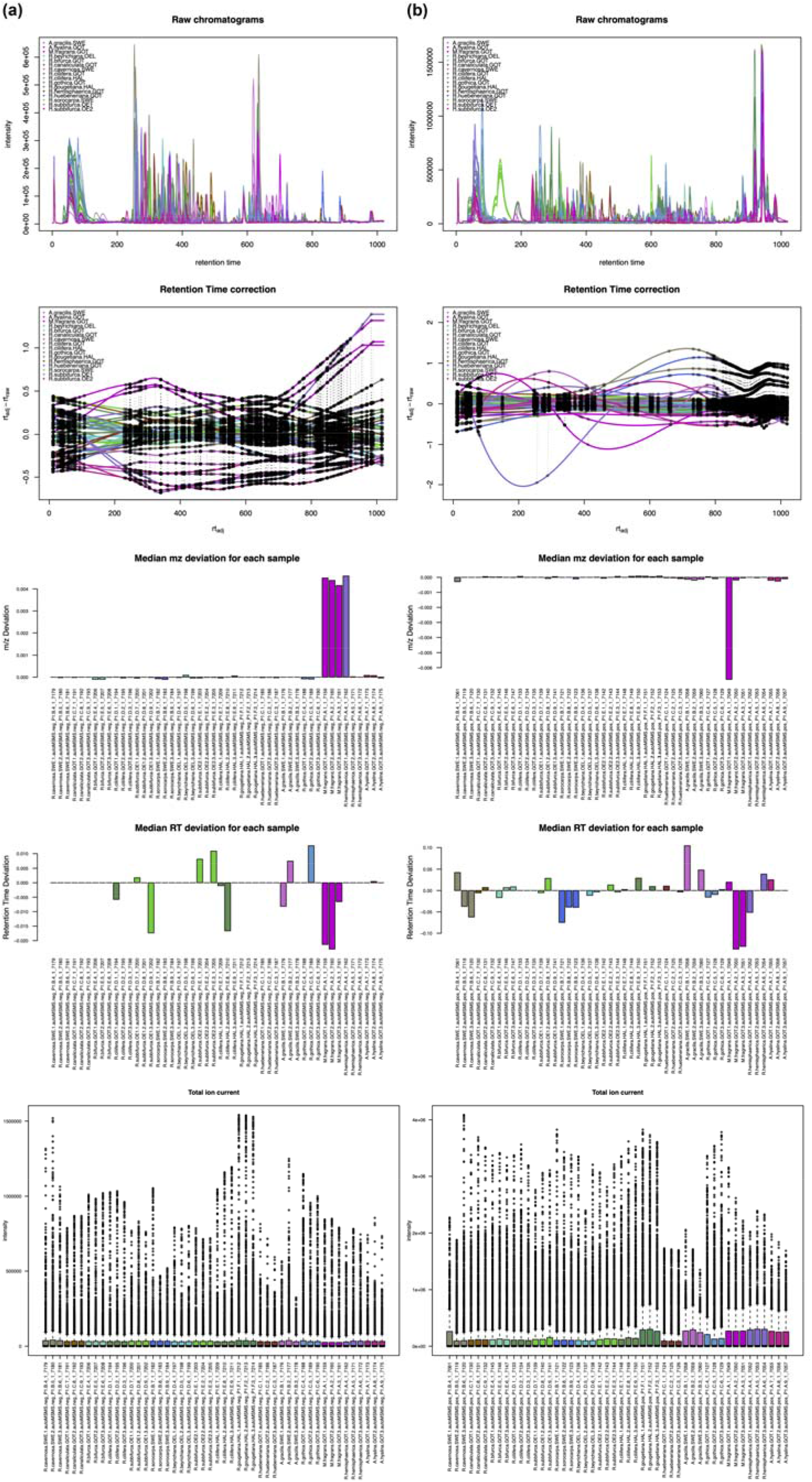
Quality Control (QC) assessment performed on the metabolomics analysis. **(a)** Negative ionisation mode. **(b)** Positive ionisation mode. Plots from top to bottom: Raw chromatograms. Retention time (RT) correction. Median mass-to-charge (mz) deviation for each sample. Median RT deviation for each sample. Total ion current (TIC) for each sample.

**Figure S2.**
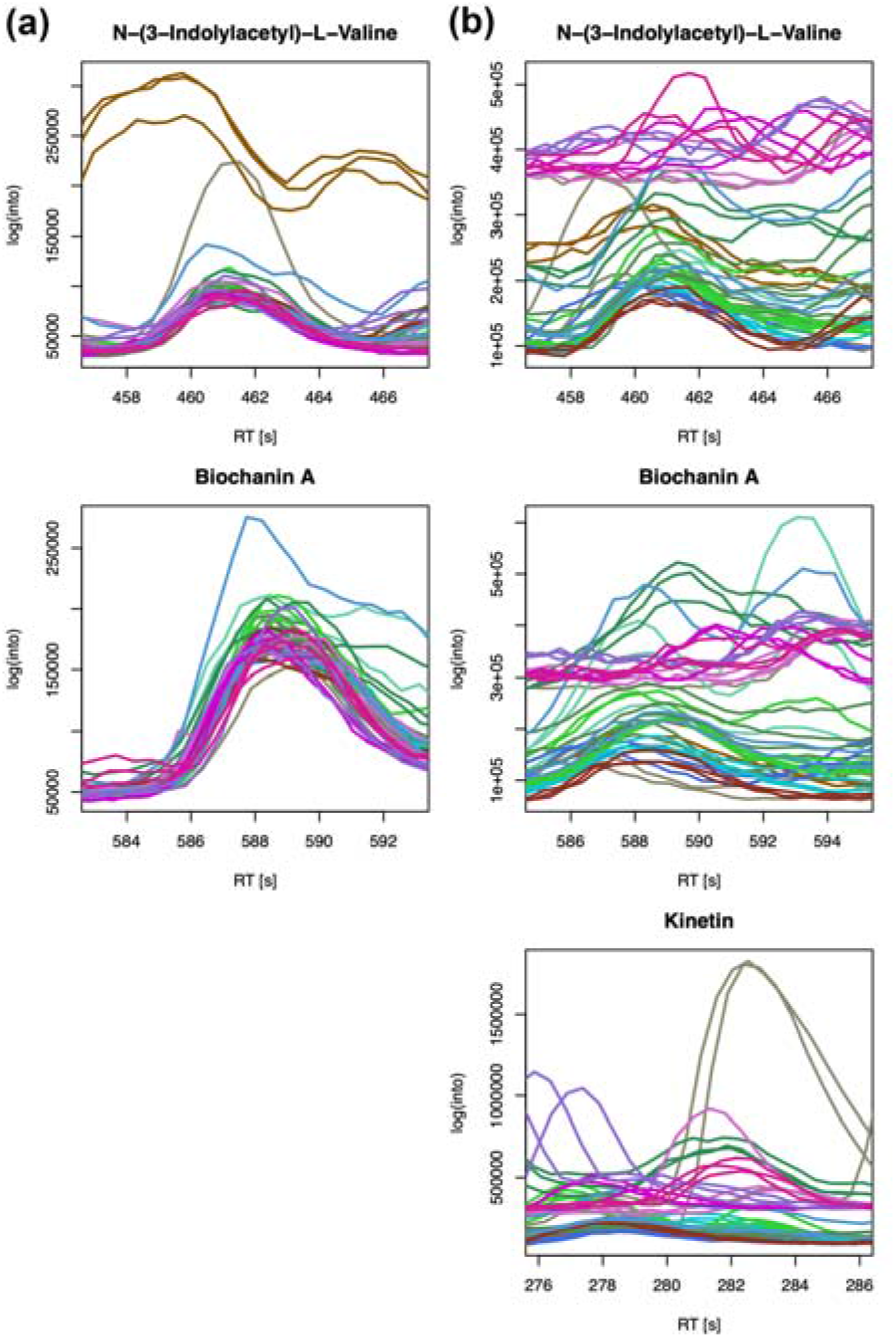
Raw chromatograms of the three internal lab standards N−(3−Indolylacetyl)−L−Valine, Biochanin A and Kinetin (only ionises in positive mode) before the alignment of XCMS. **(a)** negative ionisation mode. **(b)** positive ionization mode.

### Estimation of molecular traits for TRY

**Table S1.**
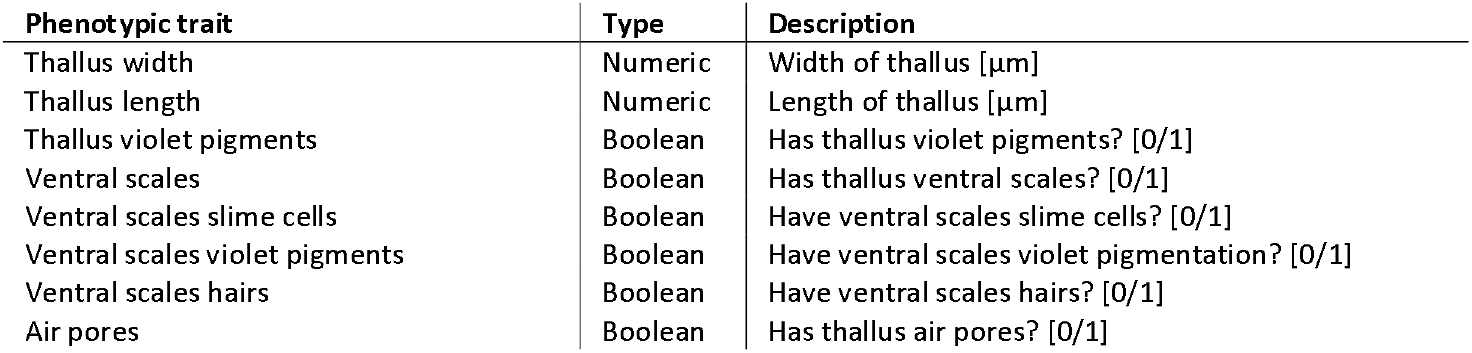
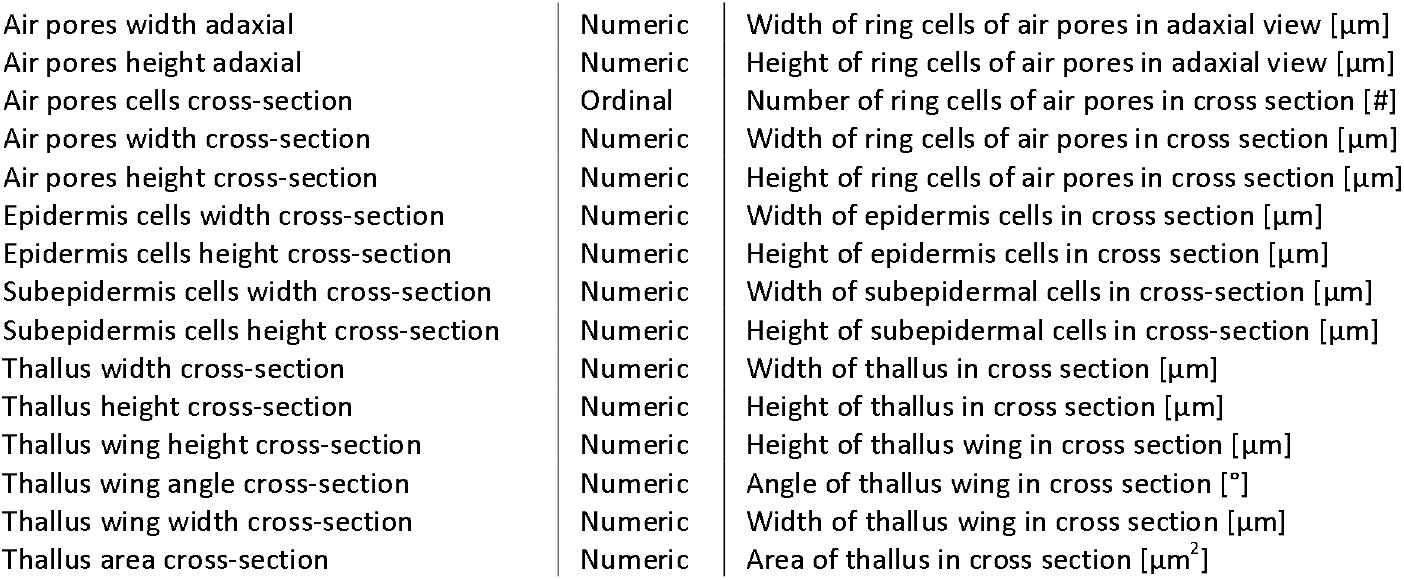
Phenotypic traits representing morphometric measurements in the investigated thallose liverworts. The column phenotypic trait contains the proposed names for the measured morphological characters. Variance, skewness and kurtosis (not shown in Table) were additionally calculated on the continuous numeric traits and included in the TRY submission.

**Table S2a.**
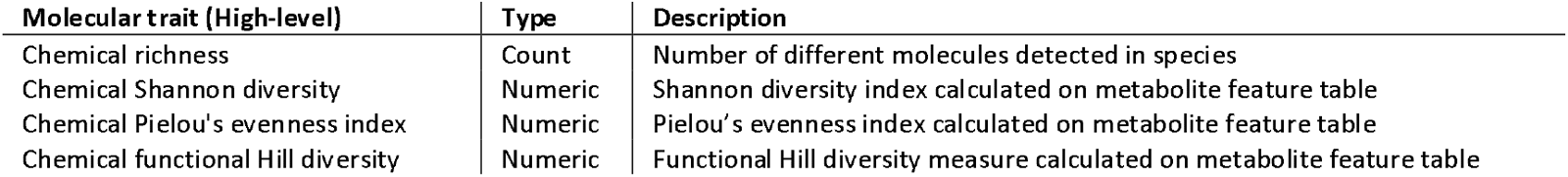
Molecular traits representing chemodiversity measures.

**Table S2b.**
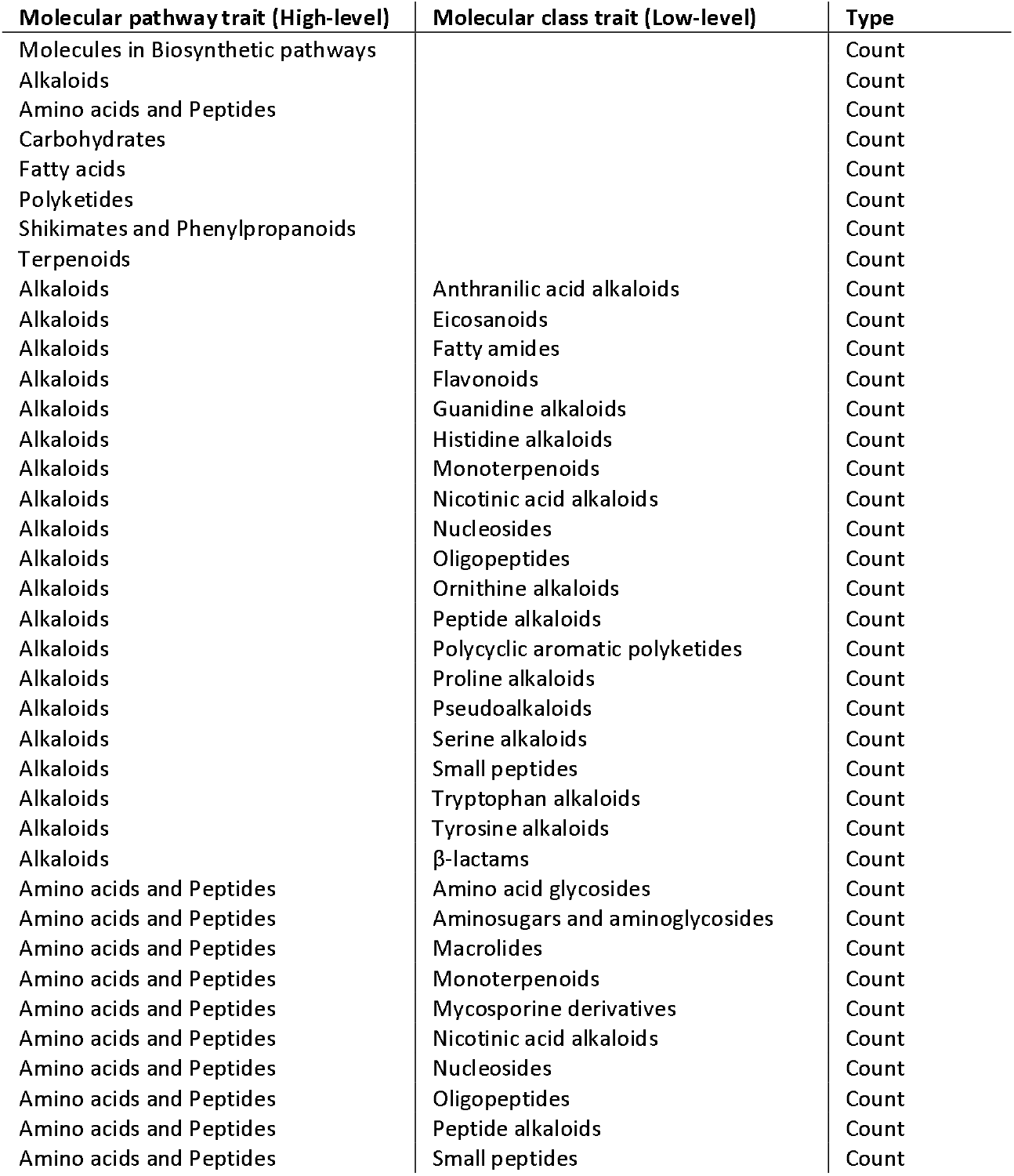
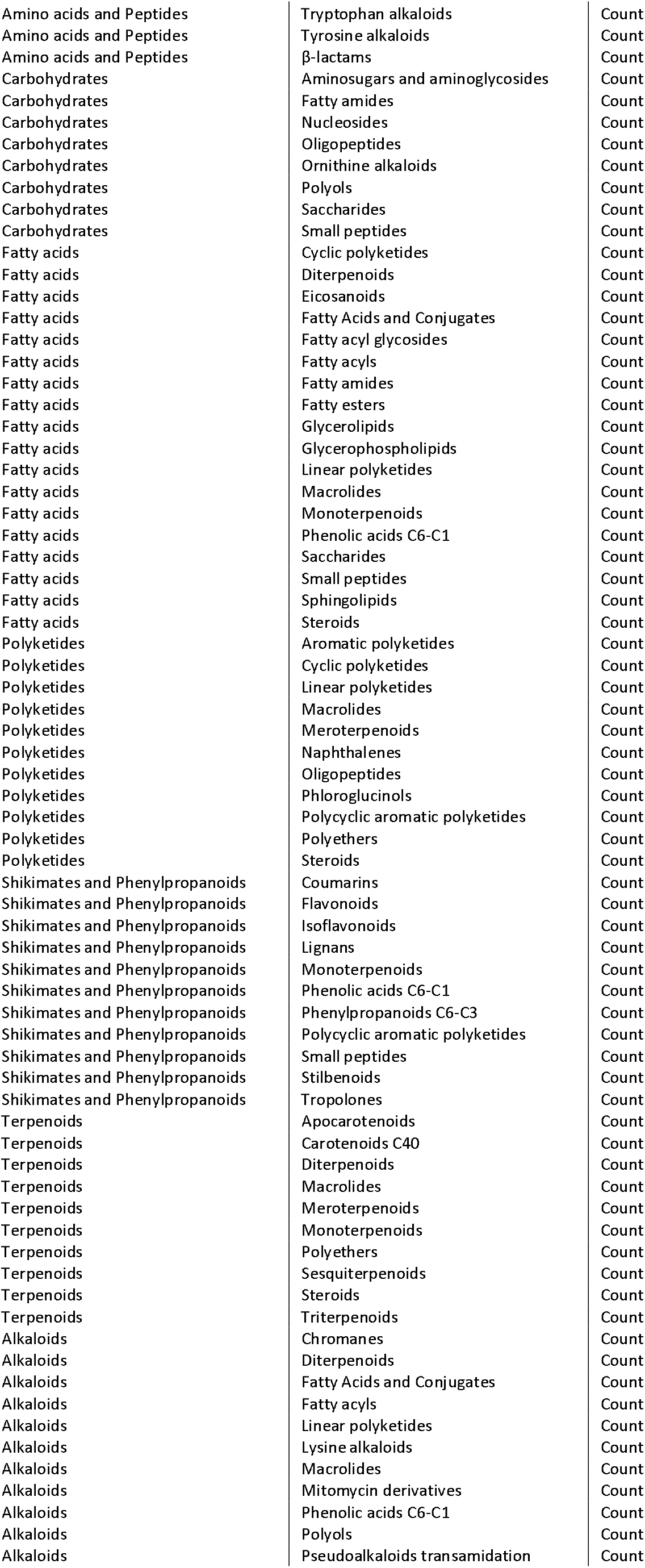
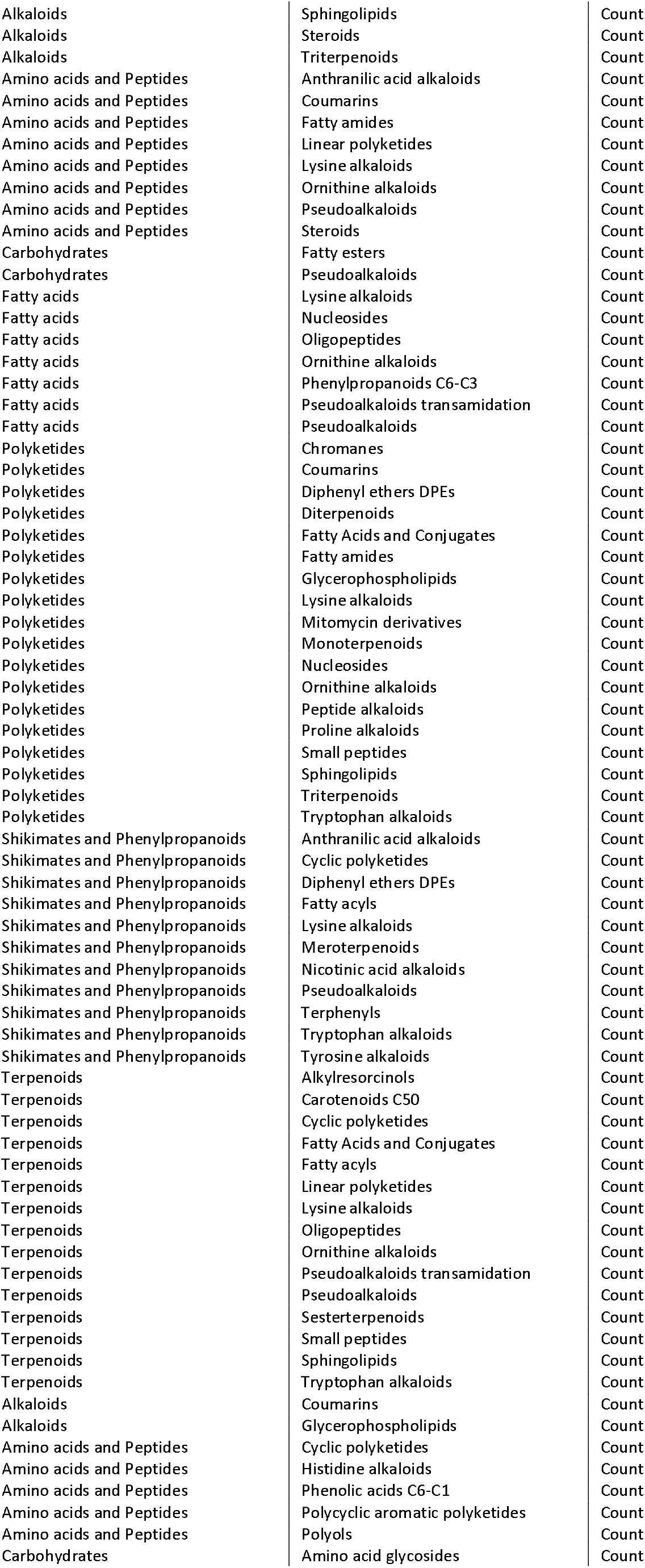
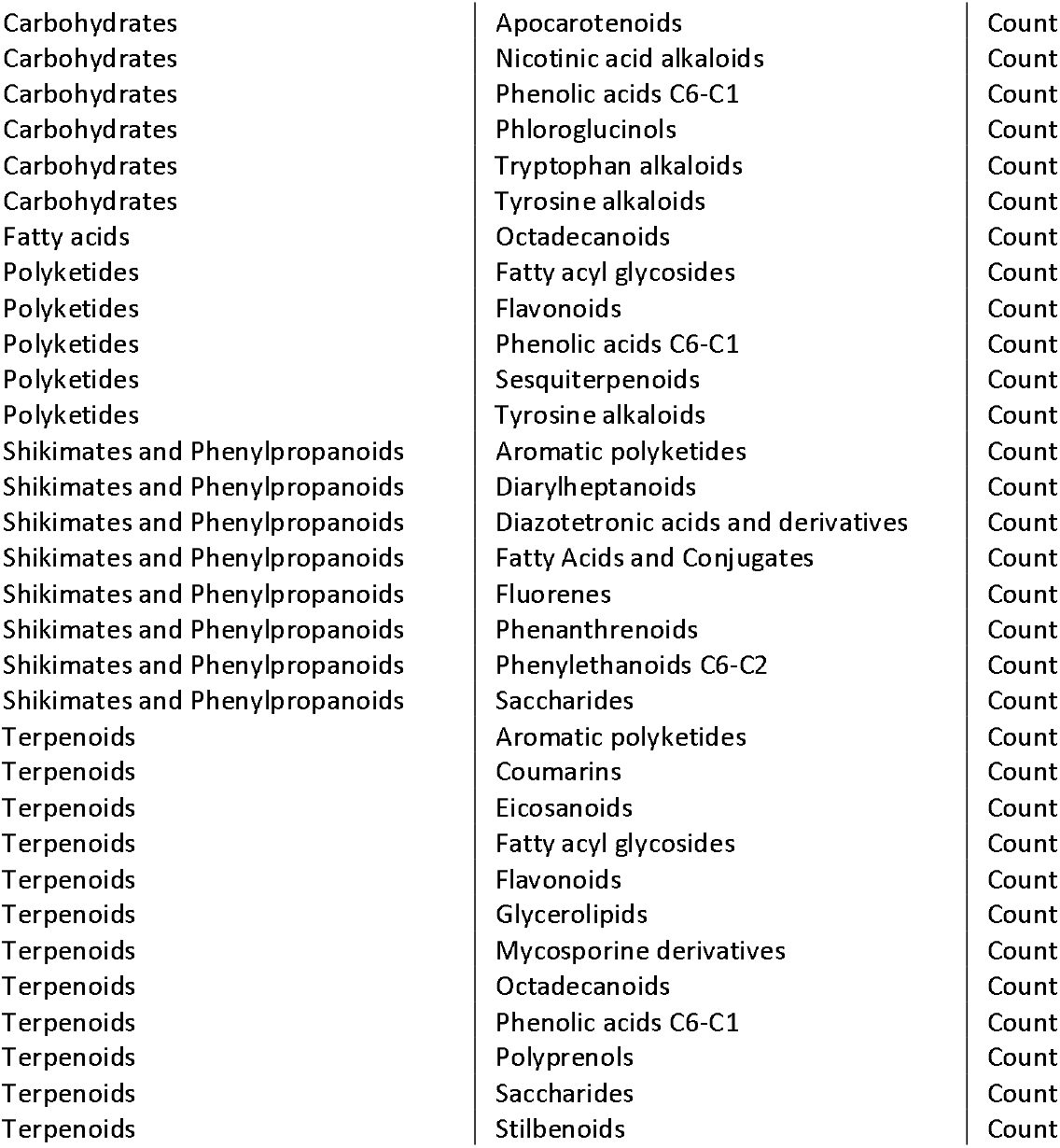
Molecular traits representing molecular pathways and classified molecules (NP-Classifier ontology). The first and second columns contain the proposed names for the high- and low-level molecular traits representing molecular pathways (high-level) and class names obtained from NP-Classifier^18^.

**Table S2c.**
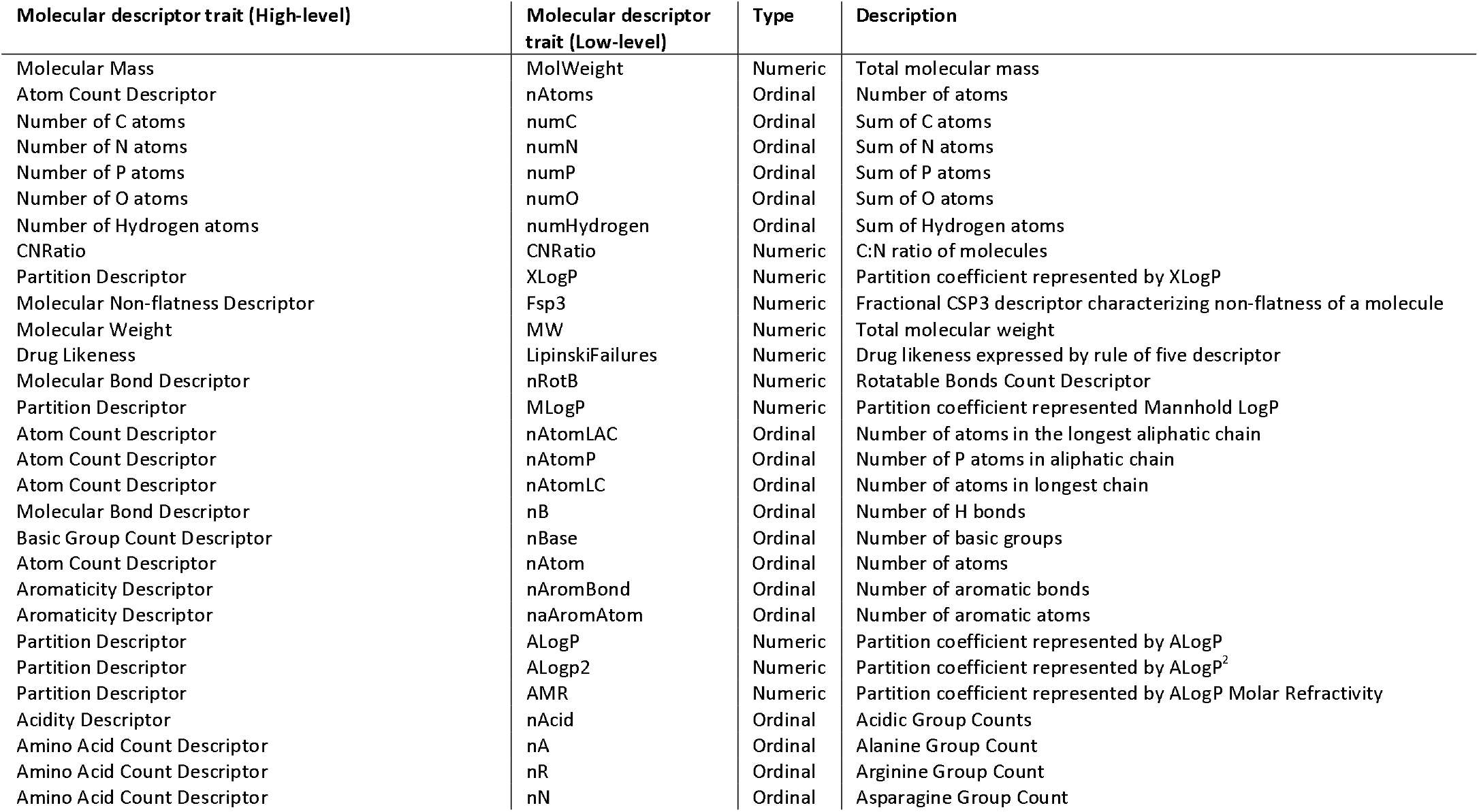
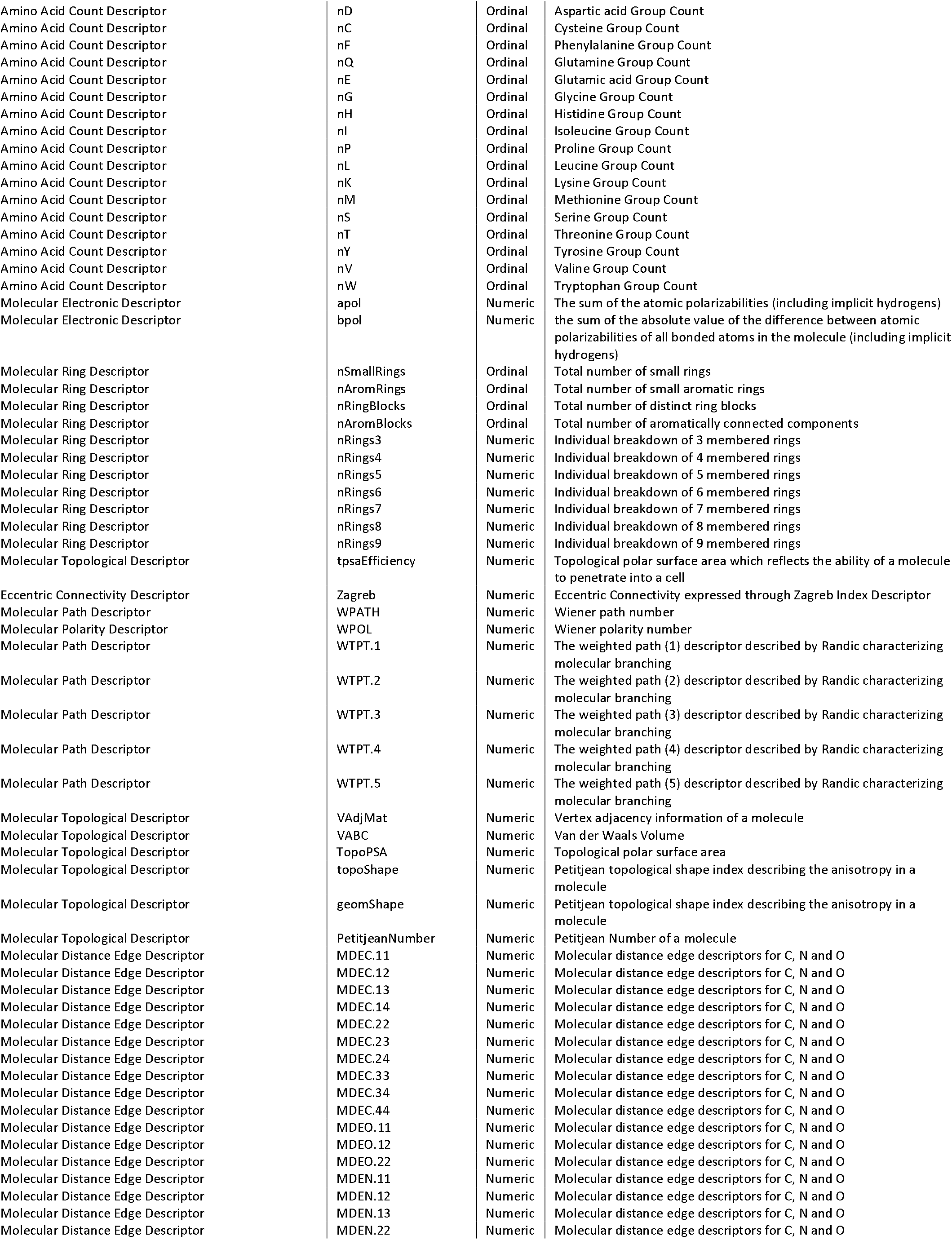
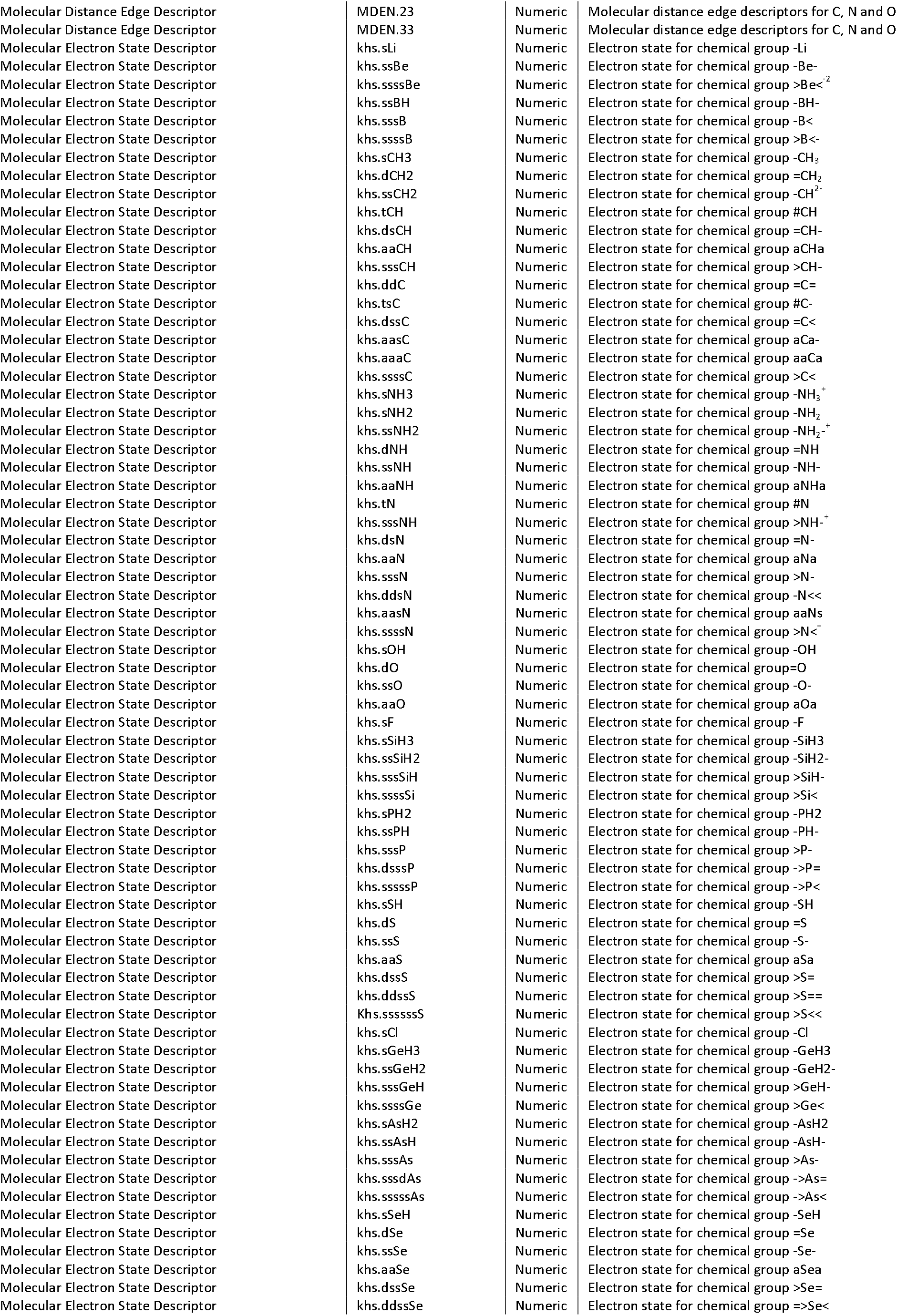
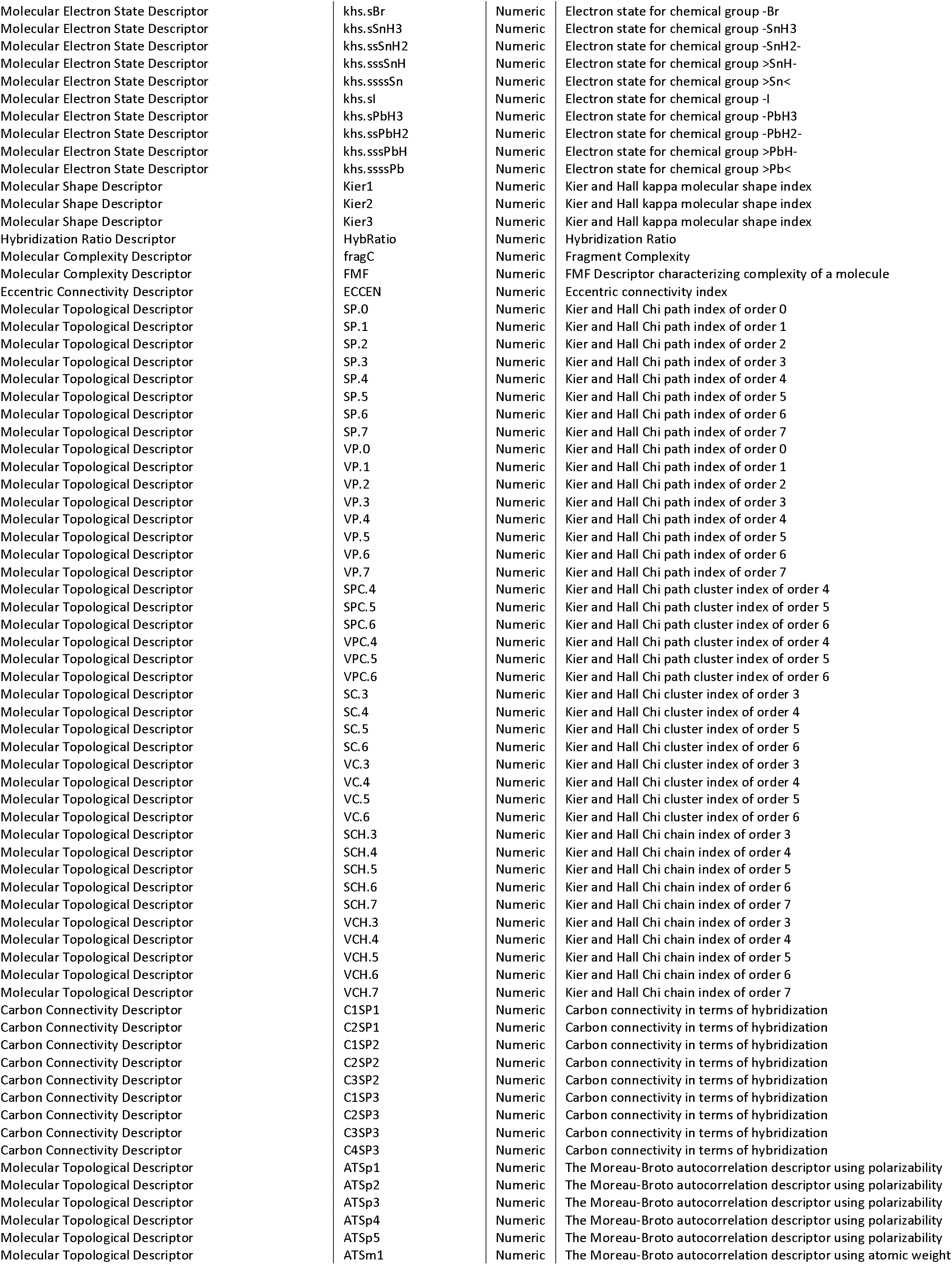
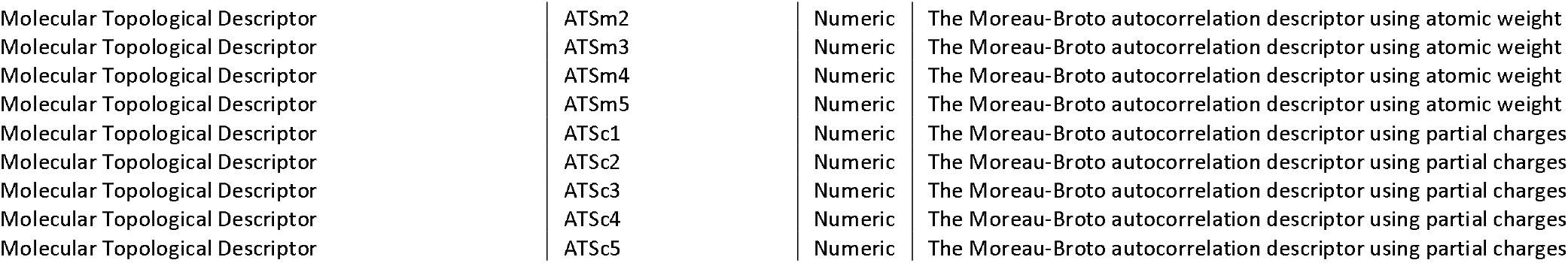
Molecular traits represented by 241 molecular descriptors. The first column contains the high-level information summarizing on mathematically similar molecular descriptors describing similar chemical functions. The second column contains the low-level information represented by the respective identifiers of the CDK nomenclature. The last column contains a description of the molecular trait.

## References

1. Stanton, D. E. & Coe, K. K. 500 million years of charted territory: functional ecological traits in bryophytes. BDE 43, (2021).

2. Díaz, S. et al. The global spectrum of plant form and function. Nature 529, 167–171 (2016).

3. Wright, I. J. et al. The worldwide leaf economics spectrum. Nature 428, 821–827 (2004).

4. Van Zuijlen, K. et al. Bryophytes of Europe Traits ( BET ) data set: A fundamental tool for ecological studies. J Vegetation Science 34, e13179 (2023).

5. Flores, J. R. et al. Dating the evolution of the complex thalloid liverworts (Marchantiopsida): total-evidence dating analysis supports a Late Silurian-Early Devonian origin and post-Mesozoic morphological stasis. New Phytologist 240, 2137–2150 (2023).

6. Hu, R. et al. Adaptive evolution of the enigmatic Takakia now facing climate change in Tibet. Cell S0092867423007365 (2023) doi:10.1016/j.cell.2023.07.003.

7. Bowman, J. L. et al. Insights into Land Plant Evolution Garnered from the Marchantia polymorpha Genome. Cell 171, 287–304.e15 (2017).

8. Walker, T. W. et al. Leaf metabolic traits reveal hidden dimensions of plant form and function. Science Advances 9, eadi4029 (2023).

9. Peters, K., Gorzolka, K., Bruelheide, H. & Neumann, S. Seasonal variation of secondary metabolites in nine different bryophytes. Ecology and Evolution 8, 9105–9117 (2018).

10. Peters, K., Poeschl, Y., Blatt-Janmaat, K. L. & Uthe, H. Ecometabolomics Studies of Bryophytes. in Bioactive Compounds in Bryophytes and Pteridophytes (ed. Murthy, H. N.) 1–43 (Springer International Publishing, 2022). doi:10.1007/978-3-030-97415-2_30-1.

11. Cornelissen, J. H. C., Lang, S. I., Soudzilovskaia, N. A. & During, H. J. Comparative Cryptogam Ecology: A Review of Bryophyte and Lichen Traits that Drive Biogeochemistry. Annals of Botany 99, 987–1001 (2007).

12. Kattge, J. et al. TRY - a global database of plant traits: TRY - A GLOBAL DATABASE OF PLANT TRAITS. Global Change Biology 17, 2905–2935 (2011).

13. Hadacek, F. Secondary Metabolites as Plant Traits: Current Assessment and Future Perspectives. Critical Reviews in Plant Sciences 21, 273–322 (2002).

14. Dührkop, K. et al. SIRIUS 4: a rapid tool for turning tandem mass spectra into metabolite structure information. Nature Methods 16, 299–302 (2019).

15. Ruttkies, C., Schymanski, E. L., Wolf, S., Hollender, J. & Neumann, S. MetFrag relaunched: incorporating strategies beyond in silico fragmentation. Journal of Cheminformatics 8, (2016).

16. Dührkop, K. et al. Systematic classification of unknown metabolites using high-resolution fragmentation mass spectra. Nat Biotechnol (2020) doi:10.1038/s41587-020-0740-8.

17. Djoumbou Feunang, Y., et al. ClassyFire: automated chemical classification with a comprehensive, computable taxonomy. J Cheminform 8, 61 (2016).

18. Gaudry, A. et al. A Sample-Centric and Knowledge-Driven Computational Framework for Natural Products Drug Discovery. https://chemrxiv.org/engage/chemrxiv/article-details/649ea085ba3e99daef499a00 (2023) doi:10.26434/chemrxiv-2023-sljbt.

19. Weininger, D. SMILES, a chemical language and information system. 1. Introduction to methodology and encoding rules. J. Chem. Inf. Comput. Sci. 28, 31–36 (1988).

20. Heller, S., McNaught, A., Stein, S., Tchekhovskoi, D. & Pletnev, I. InChI - the worldwide chemical structure identifier standard. J Cheminform 5, 7 (2013).

21. Consonni, V. Handbook of molecular descriptor.s (Wiley-VCH, 2011).

22. Mauri, A., Consonni, V. & Todeschini, R. Molecular Descriptors. in Handbook of Computational Chemistry (eds. Leszczynski, J. et al.) 2065–2093 (Springer International Publishing, 2017). doi:10.1007/978-3-319-27282-5_51.

23. Waagmeester, A. et al. Wikidata as a knowledge graph for the life sciences. eLife 9, e52614 (2020).

24. Steinbeck, C. et al. The Chemistry Development Kit (CDK): An Open-Source Java Library for Chemo- and Bioinformatics. J. Chem. Inf. Comput. Sci .43, 493–500 (2003).

25. Rutz, A. et al. The LOTUS Initiative for Open Natural Products Research: Knowledge Management through Wikidata. 78 (2021).

26. Wilkinson, M. D. et al. The FAIR Guiding Principles for scientific data management and stewardship. Scientific Data 3, 160018 (2016).

27. Pereira, H. M. et al. Essential Biodiversity Variables. Science 339, 277–278 (2013).

28. Schlick-Steiner, B. C. et al. Integrative Taxonomy: A Multisource Approach to Exploring Biodiversity. Annu. Rev. Entomol. 55, 421–438 (2010).

29. Anantaprayoon, N., Wonnapinij, P. & Kraichak, E. Integrative approaches to a revision of the liverwort in genus *Aneura* (Aneuraceae, Marchantiophyta) from Thailand. PeerJ 11, e16284 (2023).

30. Hutchinson, G. E. Concluding Remarks. Cold Spring Harbor Symposia on Quantitative Biology 22, 415–427 (1957).

31. Violle, C. et al. The return of the variance: intraspecific variability in community ecology. Trends in Ecology & Evolution 27, 244–252 (2012).

32. Lamanna, C. et al. Functional trait space and the latitudinal diversity gradient. Proc. Natl. Acad. Sci. U.S.A. 111, 13745–13750 (2014).

33. Rdusseeun, L. & Kaufman, P. Clustering by means of medoids. in Proceedings of the statistical data analysis based on the L1 norm conference, neuchatel, switzerland vol. 31 (1987).

34. Kim, H. W. et al. NPClassifier: A Deep Neural Network-Based Structural Classification Tool for Natural Products. J. Nat. Prod. 84, 2795–2807 (2021).

35. Stanstrup, J. et al. The metaRbolomics Toolbox in Bioconductor and beyond. 55 (2019).

36. Goble, C. et al. FAIR Computational Workflows. Data Intellegence 2, 108–121 (2020).

37. van Dam, N. M. & van der Meijden, E. A Role for Metabolomics in Plant Ecology. in Annual Plant Reviews Volume 43 (ed. Hall, R. D.) 87–107 (Wiley-Blackwell, 2011). doi:10.1002/9781444339956.ch4.

38. Peters, K. et al. Current Challenges in Plant Eco-Metabolomics. International Journal of Molecular Sciences 19, 1385 (2018).

39. Walker, T. W. N. et al. Functional Traits 2.0: The power of the metabolome for ecology. Journal of Ecology 110, 4–20 (2022).

40. Peters, K. & König-Ries, B. Reference bioimaging to assess the phenotypic trait diversity of bryophytes within the family Scapaniaceae. Sci Data 9, 598 (2022).

41. Holmgren, P. K. & Holmgren, N. H. Index Herbariorum. Taxon 40, 687–692 (1991).

42. Nelson, G., Paul, D., Riccardi, G. & Mast, A. Five task clusters that enable efficient and effective digitization of biological collections. ZK 209, 19–45 (2012).

43. Schindelin, J., et al. Fiji: an open-source platform for biological-image analysis. Nat Methods 9, 676–682 (2012).

44. Arzt, M., et al. LABKIT: Labeling and Segmentation Toolkit for Big Image Data. Front. Comput. Sci. 4, 777728 (2022).

45. Weigert, M., Schmidt, U., Haase, R., Sugawara, K. & Myers, G. Star-convex Polyhedra for 3D Object Detection and Segmentation in Microscopy. in 2020 IEEE Winter Conference on Applications of Computer Vision (WACV) 3655–3662 (IEEE, 2020). doi:10.1109/WACV45572.2020.9093435.

46. Enquist, B. J. et al. Scaling from Traits to Ecosystems. in Advances in Ecological Research vol. 52 249–318 (Elsevier, 2015).

47. Zhang, L., Bai, W., Zhang, Y., Lambers, H. & Zhang, W. Ecosystem stability is determined by plant defence functional traits and population stability under mowing in a semi-arid temperate steppe. Functional Ecology 37, 2413–2424 (2023).

48. Stech, M. & Quandt, D. 20,000 species and five key markers: The status of molecular bryophyte phylogenetics. Phytotaxa 9, 196 (2014).

49. Edgar, R. C. MUSCLE: multiple sequence alignment with high accuracy and high throughput. Nucleic Acids Research 32, 1792–1797 (2004).

50. Burgin, J. et al. The European Nucleotide Archive in 2022. Nucleic Acids Research 51, D121–D125 (2023).

51. Stamatakis, A. RAxML version 8: a tool for phylogenetic analysis and post-analysis of large phylogenies. Bioinformatics 30, 1312–1313 (2014).

52. Benson, D. A. et al. GenBank. Nucleic Acids Research 41, D36–D42 (2012).

53. Thiele, K. The Holy Grail of the Perfect Character: the Cladistic Treatment of Morphometric Data. Cladistics 9, 275–304 (1993).

54. Garcia-Cruz, J. & Sosa, V. Coding Quantitative Character Data for Phylogenetic Analysis: A Comparison of Five Methods. issn: 0363-6445 31, 302–309 (2006).

55. Jordon-Thaden, I. E., Chanderbali, A. S., Gitzendanner, M. A. & Soltis, D. E. Modified CTAB and TRIzol protocols improve RNA extraction from chemically complex Embryophyta. Applications in Plant Sciences 3, 1400105 (2015).

56. Böttcher, C. et al. The Multifunctional Enzyme CYP71B15 (PHYTOALEXIN DEFICIENT3) Converts Cysteine-Indole-3-Acetonitrile to Camalexin in the Indole-3-Acetonitrile Metabolic Network of Arabidopsis thaliana. THE PLANT CELL ONLINE 21, 1830–1845 (2009).

57. Blatt-Janmaat, K. L. et al. Impact of in vitro hormone treatments on the bibenzyl production of Radula complanata. Botany cjb-2022-0048 (2022) doi:10.1139/cjb-2022-0048.

58. Peters, K., Blatt-Janmaat, K. L., Tkach, N., Van Dam, N. M. & Neumann, S. Untargeted Metabolomics for Integrative Taxonomy: Metabolomics, DNA Marker-Based Sequencing, and Phenotype Bioimaging. Plants 12, 881 (2023).

59. Chambers, M. C. et al. A cross-platform toolkit for mass spectrometry and proteomics. Nature Biotechnology 30, 918–920 (2012).

60. Haug, K. et al. MetaboLights—an open-access general-purpose repository for metabolomics studies and associated meta-data. Nucleic Acids Research 41, D781–D786 (2013).

61. Spicer, R. A., Salek, R. & Steinbeck, C. Compliance with minimum information guidelines in public metabolomics repositories. Scientific Data 4, 170137 (2017).

62. Smith, C. A., Want, E. J., O’Maille, G., Abagyan, R. & Siuzdak, G. XCMS: Processing Mass Spectrometry Data for Metabolite Profiling Using Nonlinear Peak Alignment, Matching, and Identification. Analytical Chemistry 78, 779–787 (2006).

63. Hoffmann, M. A. et al. High-confidence structural annotation of metabolites absent from spectral libraries. Nat Biotechnol 40, 411–421 (2022).

64. Voicu, A., Duteanu, N., Voicu, M., Vlad, D. & Dumitrascu, V. The rcdk and cluster R packages applied to drug candidate selection. J Cheminform 12, 3 (2020).

65. Peters et al. Chemical Diversity and Classification of Secondary Metabolites in Nine Bryophyte Species. Metabolites 9, 222 (2019).

66. Grau, J., Grosse, I. & Keilwagen, J. PRROC: computing and visualizing precision-recall and receiver operating characteristic curves in R. Bioinformatics 31, 2595–2597 (2015).

67. Robin, X. et al. pROC: an open-source package for R and S+ to analyze and compare ROC curves. BMC Bioinformatics 12, 77 (2011).

68. Fawcett, T. An introduction to ROC analysis. Pattern Recognition Letters 27, 861–874 (2006).

69. Tharwat, A. Classification assessment methods. ACI ahead-of-print, (2020).

