## Supplemental Material for "Estimating essential phenotypic and molecular traits from integrative biodiversity data"

**Affiliations**

### Supplemental Material

***Technical validation of metabolomics data***

The 48 samples were run in randomized injection order in one instrument run to avoid batch effects. Diagnostic plots of XCMS were employed to validate instrument performance and to detect shifts between the instrument runs. First, raw chromatograms were visually inspected for any shifts in intensity (Fig. S1, first row). Next, retention time (RT) correction made by XCMS on the samples was assessed (Fig. S1, second row) followed by mass-to-charge deviation (m/z) (in ppm) (Fig. S1, third row) and RT deviation (in seconds) (Fig. S1, fourth row). Lastly, total ion current (TIC) was determined and compared for the samples (Fig. S1, last row). In summary, we found maximum retention time deviations within 2 seconds – which is well within limits of the analytical setup used. The determined maximum mass-to-charge deviations of 4 ppm (negative ion mode) and 6 ppm (positive mode) were within instrument specification as well. For the majority of samples, the deviations were even less than 1 ppm (Fig. S1, third row). Deviations in mass-to-charge and retention times were corrected by XCMS. We conclude that there are no significant shifts in the instrument run and that the corrections made by XCMS are validated for the parameters used for peak detection.

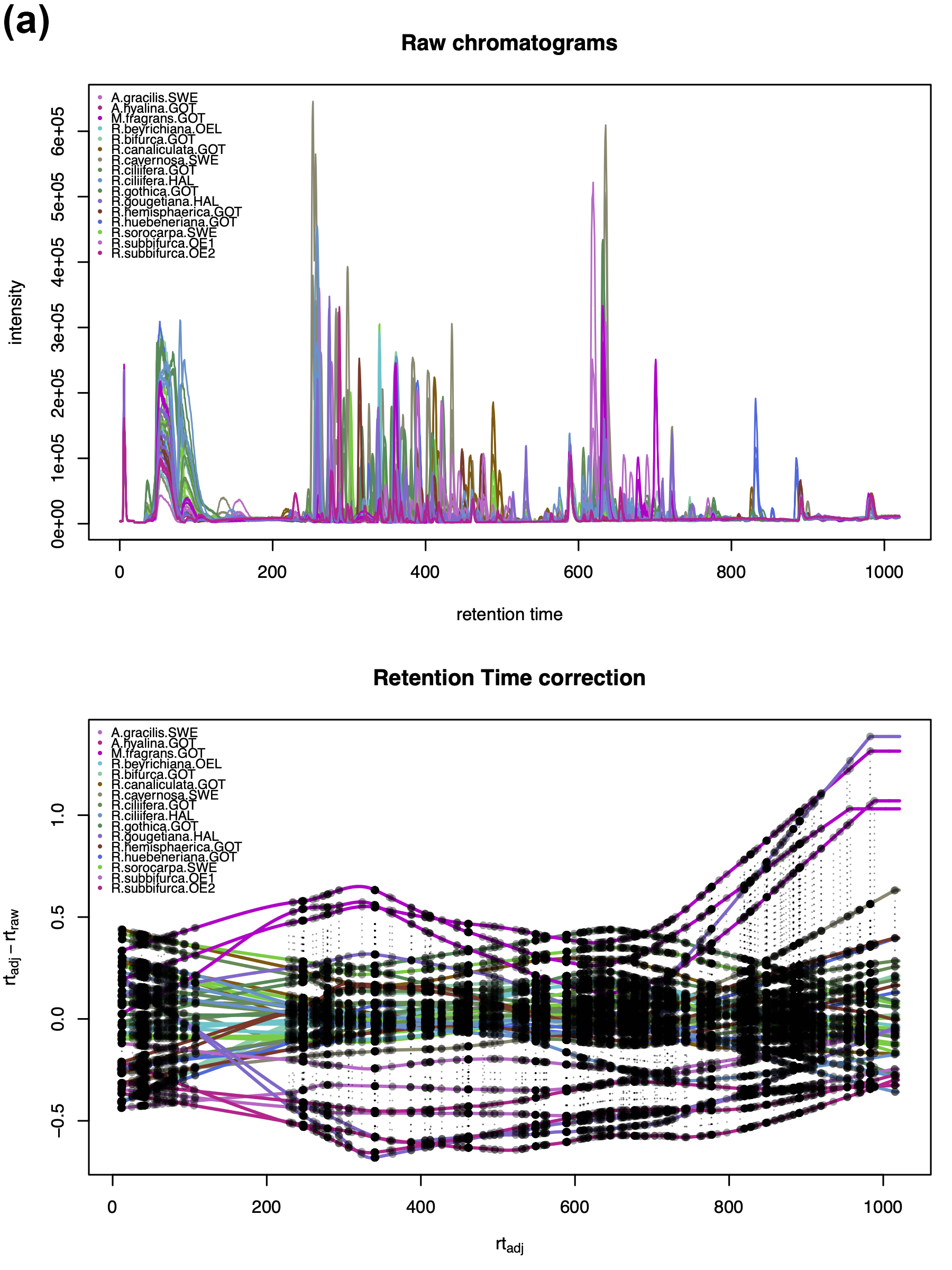

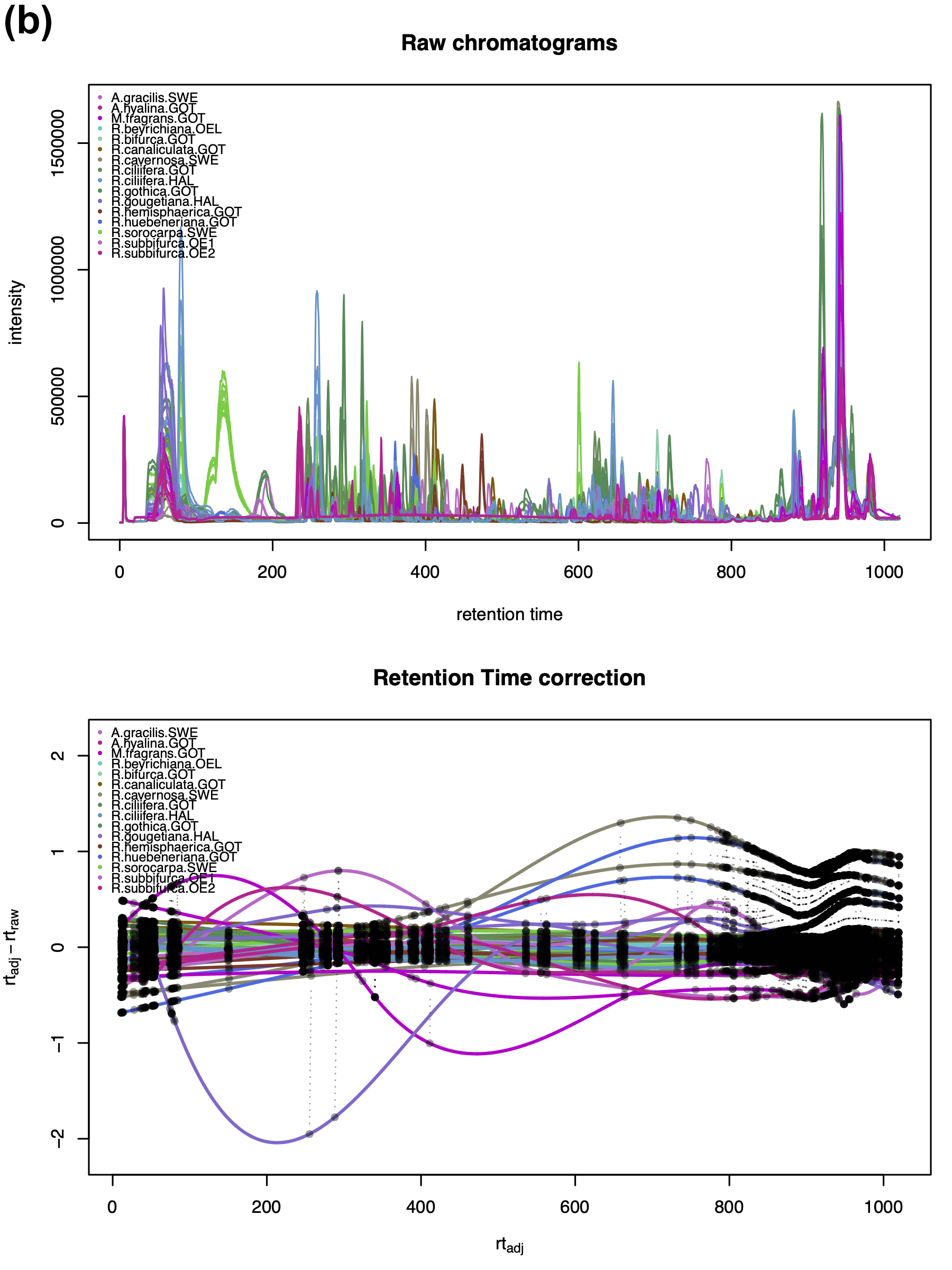

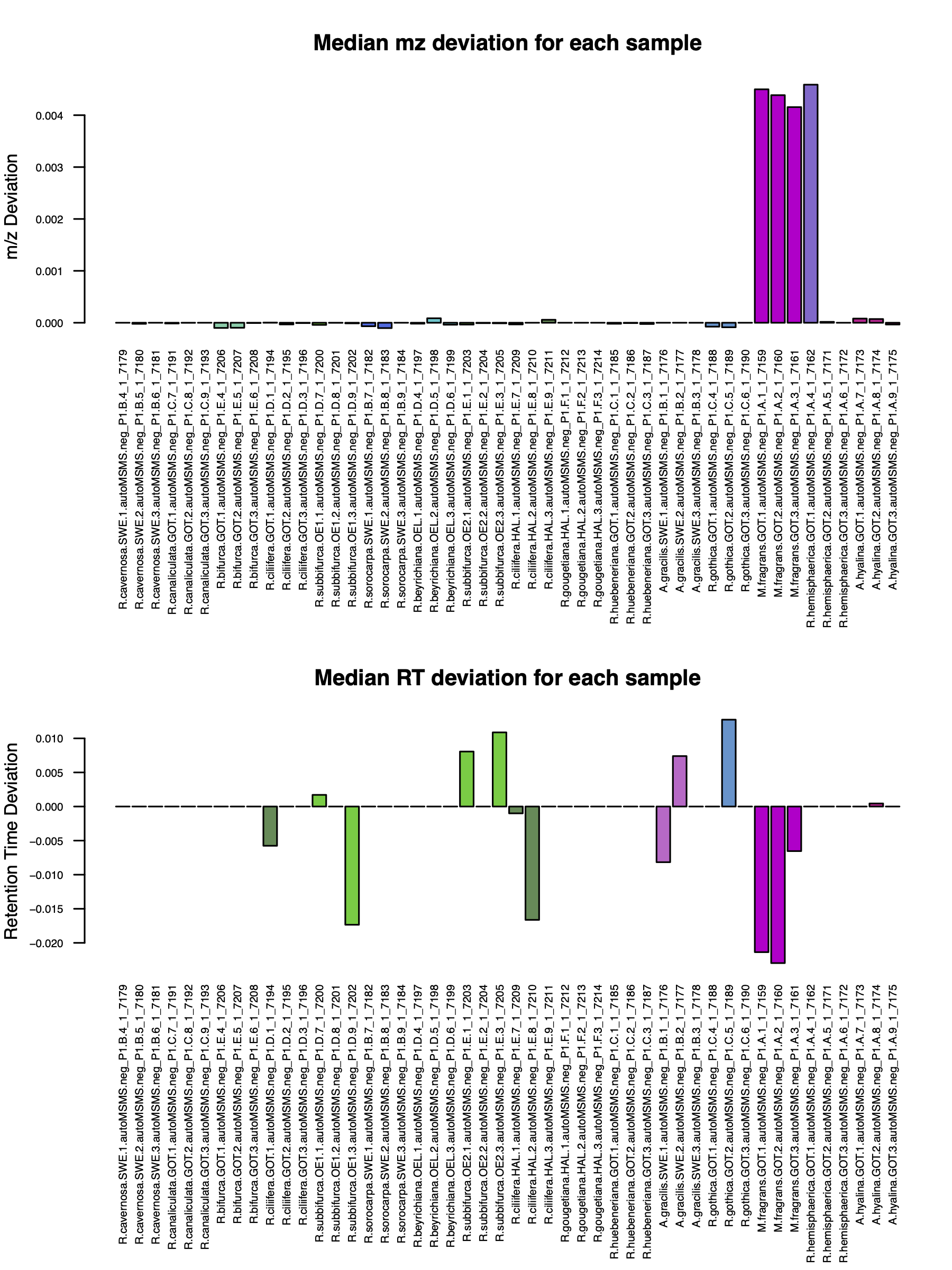

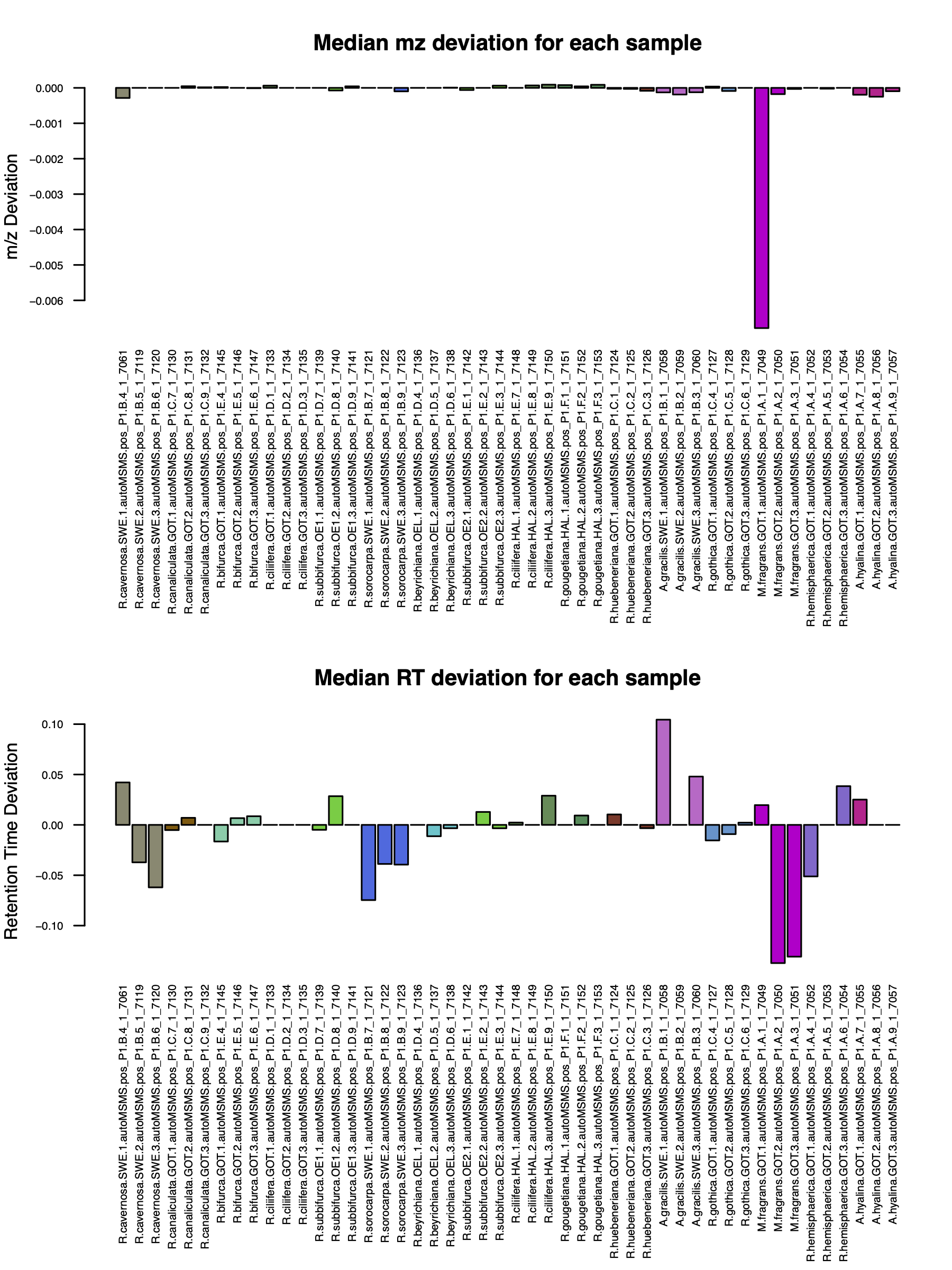

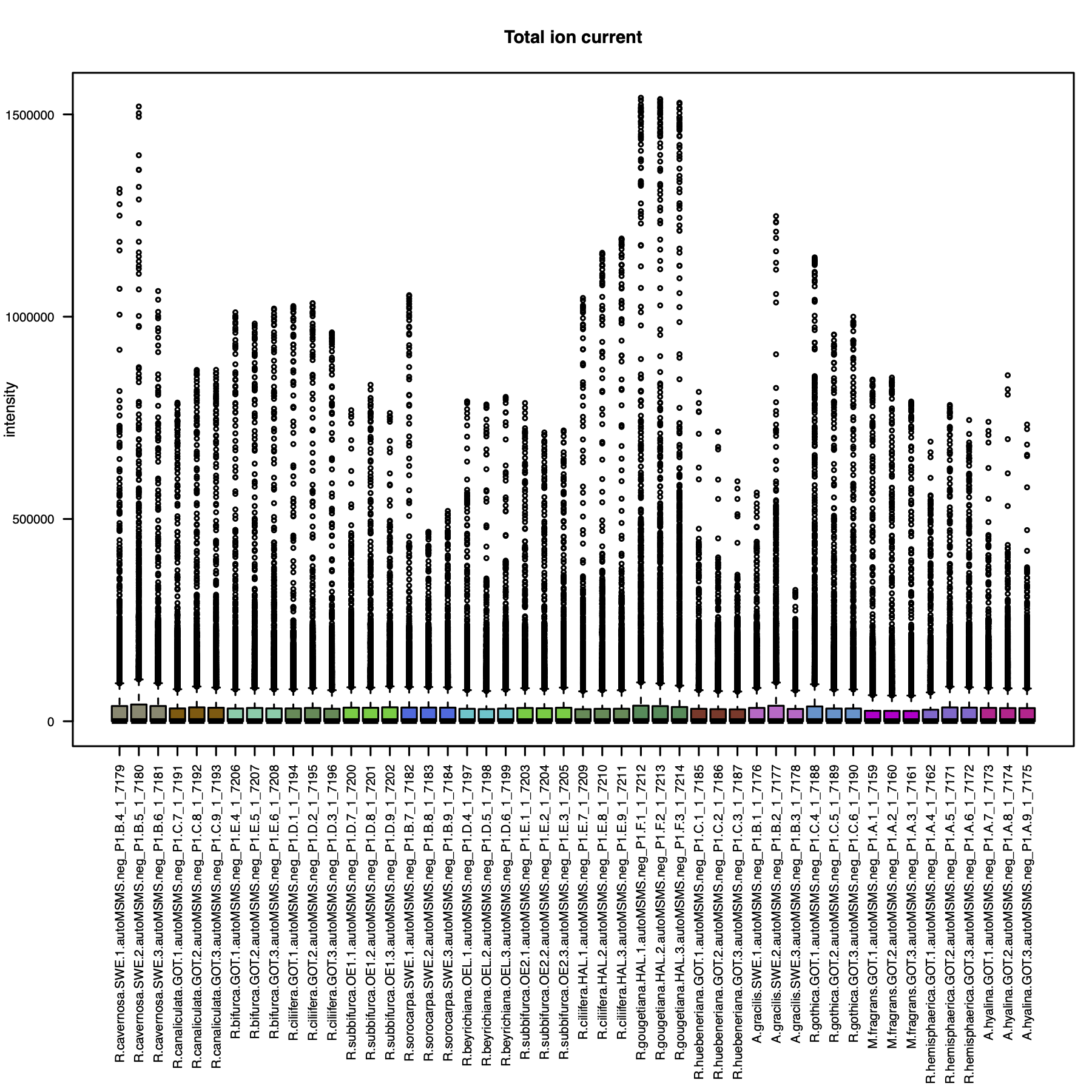

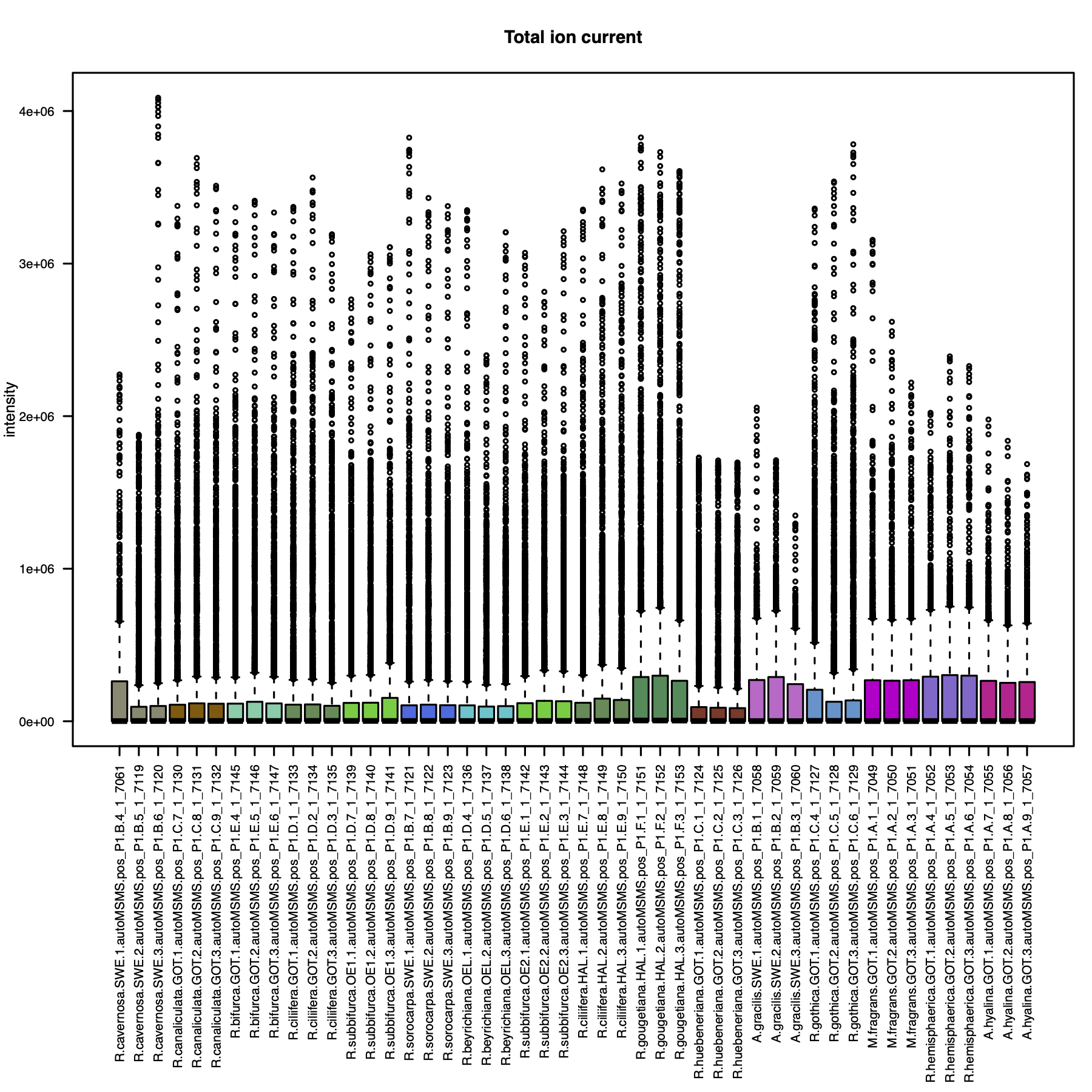

**Figure S1.** Quality Control (QC) assessment performed on the metabolomics analysis. **(a)** Negative ionisation mode. **(b)** Positive ionisation mode. Plots from top to bottom: Raw chromatograms. Retention time (RT) correction. Median mass-to-charge (mz) deviation for each sample. Median RT deviation for each sample. Total ion current (TIC) for each sample.

In addition, the intensities of three internal lab standards that were spiked into all samples were also investigated (Fig. S2). We found that the variation for the internal lab standard were within the typical range of 10-15%^52^.

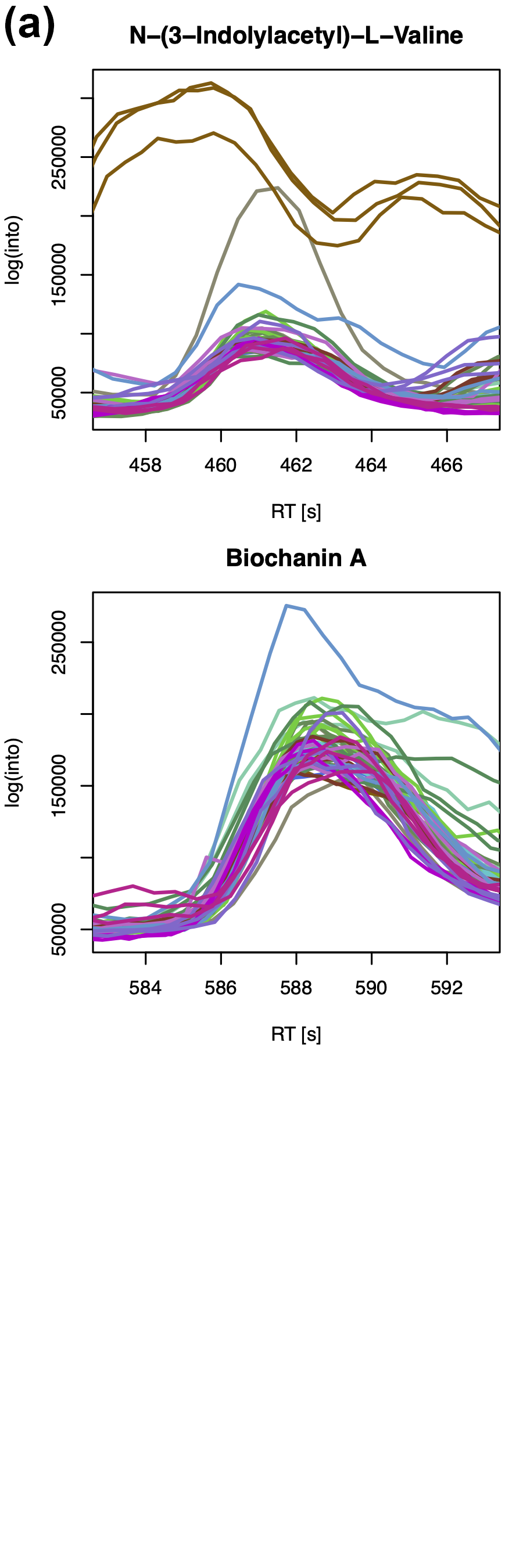

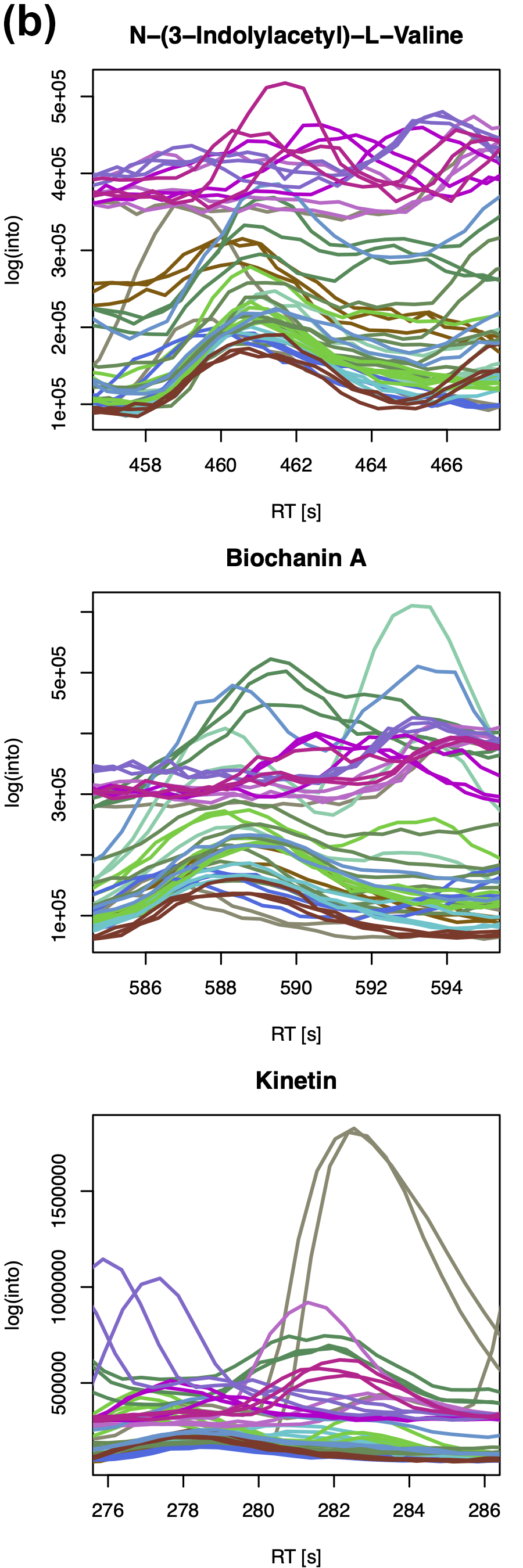

**Figure S2.** Raw chromatograms of the three internal lab standards N−(3−Indolylacetyl)−L−Valine, Biochanin A and Kinetin (only ionises in positive mode) before the alignment of XCMS. **(a)** negative ionisation mode. **(b)** positive ionization mode.

***Estimation of morphometric traits***

**Table S1.** Phenotypic traits representing morphometric measurements in the investigated thallose liverworts. The column phenotypic trait contains the proposed names for the measured morphological characters. Variance, skewness and kurtosis (not shown in Table) were additionally calculated on the continuous numeric traits and included in the TRY submission.

| **Phenotypic trait** | **Type** | **Description** |
| --- | --- | --- |
| Thallus width | Numeric | Width of thallus [µm] |
| Thallus length | Numeric | Length of thallus [µm] |
| Thallus violet pigments | Boolean | Has thallus violet pigments? [0/1] |
| Ventral scales | Boolean | Has thallus ventral scales? [0/1] |
| Ventral scales slime cells | Boolean | Have ventral scales slime cells? [0/1] |
| Ventral scales violet pigments | Boolean | Have ventral scales violet pigmentation? [0/1] |
| Ventral scales hairs | Boolean | Have ventral scales hairs? [0/1] |
| Air pores | Boolean | Has thallus air pores? [0/1] |
| Air pores width adaxial | Numeric | Width of ring cells of air pores in adaxial view [µm] |
| Air pores height adaxial | Numeric | Height of ring cells of air pores in adaxial view [µm] |
| Air pores cells cross-section | Ordinal | Number of ring cells of air pores in cross section [#] |
| Air pores width cross-section | Numeric | Width of ring cells of air pores in cross section [µm] |
| Air pores height cross-section | Numeric | Height of ring cells of air pores in cross section [µm] |
| Epidermis cells width cross-section | Numeric | Width of epidermis cells in cross section [µm] |
| Epidermis cells height cross-section | Numeric | Height of epidermis cells in cross section [µm] |
| Subepidermis cells width cross-section | Numeric | Width of subepidermal cells in cross-section [µm] |
| Subepidermis cells height cross-section | Numeric | Height of subepidermal cells in cross-section [µm] |
| Thallus width cross-section | Numeric | Width of thallus in cross section [µm] |
| Thallus height cross-section | Numeric | Height of thallus in cross section [µm] |
| Thallus wing height cross-section | Numeric | Height of thallus wing in cross section [µm] |
| Thallus wing angle cross-section | Numeric | Angle of thallus wing in cross section [°] |
| Thallus wing width cross-section | Numeric | Width of thallus wing in cross section [µm] |
| Thallus area cross-section | Numeric | Area of thallus in cross section [µm^2^] |

***Estimation of molecular traits for TRY***

**Table S2a.** Molecular traits representing chemodiversity measures.

| **Molecular trait (High-level)** | **Type** | **Description** |
| --- | --- | --- |
| Chemical richness | Count | Number of different molecules detected in species |
| Chemical Shannon diversity | Numeric | Shannon diversity index calculated on metabolite feature table |
| Chemical Pielou's evenness index | Numeric | Pielou’s evenness index calculated on metabolite feature table |
| Chemical functional Hill diversity | Numeric | Functional Hill diversity measure calculated on metabolite feature table |

**Table S2b.** Molecular traits representing molecular pathways and classified molecules (NP-Classifier ontology). The first and second columns contain the proposed names for the high- and low-level molecular traits representing molecular pathways (high-level) and class names obtained from NP-Classifier^18^.

| **Molecular pathway trait (High-level)** | **Molecular class trait (Low-level)** | **Type** |
| --- | --- | --- |
| Molecules in Biosynthetic pathways |  | Count |
| Alkaloids |  | Count |
| Amino acids and Peptides |  | Count |
| Carbohydrates |  | Count |
| Fatty acids |  | Count |
| Polyketides |  | Count |
| Shikimates and Phenylpropanoids |  | Count |
| Terpenoids |  | Count |
| Alkaloids | Anthranilic acid alkaloids | Count |
| Alkaloids | Eicosanoids | Count |
| Alkaloids | Fatty amides | Count |
| Alkaloids | Flavonoids | Count |
| Alkaloids | Guanidine alkaloids | Count |
| Alkaloids | Histidine alkaloids | Count |
| Alkaloids | Monoterpenoids | Count |
| Alkaloids | Nicotinic acid alkaloids | Count |
| Alkaloids | Nucleosides | Count |
| Alkaloids | Oligopeptides | Count |
| Alkaloids | Ornithine alkaloids | Count |
| Alkaloids | Peptide alkaloids | Count |
| Alkaloids | Polycyclic aromatic polyketides | Count |
| Alkaloids | Proline alkaloids | Count |
| Alkaloids | Pseudoalkaloids | Count |
| Alkaloids | Serine alkaloids | Count |
| Alkaloids | Small peptides | Count |
| Alkaloids | Tryptophan alkaloids | Count |
| Alkaloids | Tyrosine alkaloids | Count |
| Alkaloids | β-lactams | Count |
| Amino acids and Peptides | Amino acid glycosides | Count |
| Amino acids and Peptides | Aminosugars and aminoglycosides | Count |
| Amino acids and Peptides | Macrolides | Count |
| Amino acids and Peptides | Monoterpenoids | Count |
| Amino acids and Peptides | Mycosporine derivatives | Count |
| Amino acids and Peptides | Nicotinic acid alkaloids | Count |
| Amino acids and Peptides | Nucleosides | Count |
| Amino acids and Peptides | Oligopeptides | Count |
| Amino acids and Peptides | Peptide alkaloids | Count |
| Amino acids and Peptides | Small peptides | Count |
| Amino acids and Peptides | Tryptophan alkaloids | Count |
| Amino acids and Peptides | Tyrosine alkaloids | Count |
| Amino acids and Peptides | β-lactams | Count |
| Carbohydrates | Aminosugars and aminoglycosides | Count |
| Carbohydrates | Fatty amides | Count |
| Carbohydrates | Nucleosides | Count |
| Carbohydrates | Oligopeptides | Count |
| Carbohydrates | Ornithine alkaloids | Count |
| Carbohydrates | Polyols | Count |
| Carbohydrates | Saccharides | Count |
| Carbohydrates | Small peptides | Count |
| Fatty acids | Cyclic polyketides | Count |
| Fatty acids | Diterpenoids | Count |
| Fatty acids | Eicosanoids | Count |
| Fatty acids | Fatty Acids and Conjugates | Count |
| Fatty acids | Fatty acyl glycosides | Count |
| Fatty acids | Fatty acyls | Count |
| Fatty acids | Fatty amides | Count |
| Fatty acids | Fatty esters | Count |
| Fatty acids | Glycerolipids | Count |
| Fatty acids | Glycerophospholipids | Count |
| Fatty acids | Linear polyketides | Count |
| Fatty acids | Macrolides | Count |
| Fatty acids | Monoterpenoids | Count |
| Fatty acids | Phenolic acids C6-C1 | Count |
| Fatty acids | Saccharides | Count |
| Fatty acids | Small peptides | Count |
| Fatty acids | Sphingolipids | Count |
| Fatty acids | Steroids | Count |
| Polyketides | Aromatic polyketides | Count |
| Polyketides | Cyclic polyketides | Count |
| Polyketides | Linear polyketides | Count |
| Polyketides | Macrolides | Count |
| Polyketides | Meroterpenoids | Count |
| Polyketides | Naphthalenes | Count |
| Polyketides | Oligopeptides | Count |
| Polyketides | Phloroglucinols | Count |
| Polyketides | Polycyclic aromatic polyketides | Count |
| Polyketides | Polyethers | Count |
| Polyketides | Steroids | Count |
| Shikimates and Phenylpropanoids | Coumarins | Count |
| Shikimates and Phenylpropanoids | Flavonoids | Count |
| Shikimates and Phenylpropanoids | Isoflavonoids | Count |
| Shikimates and Phenylpropanoids | Lignans | Count |
| Shikimates and Phenylpropanoids | Monoterpenoids | Count |
| Shikimates and Phenylpropanoids | Phenolic acids C6-C1 | Count |
| Shikimates and Phenylpropanoids | Phenylpropanoids C6-C3 | Count |
| Shikimates and Phenylpropanoids | Polycyclic aromatic polyketides | Count |
| Shikimates and Phenylpropanoids | Small peptides | Count |
| Shikimates and Phenylpropanoids | Stilbenoids | Count |
| Shikimates and Phenylpropanoids | Tropolones | Count |
| Terpenoids | Apocarotenoids | Count |
| Terpenoids | Carotenoids C40 | Count |
| Terpenoids | Diterpenoids | Count |
| Terpenoids | Macrolides | Count |
| Terpenoids | Meroterpenoids | Count |
| Terpenoids | Monoterpenoids | Count |
| Terpenoids | Polyethers | Count |
| Terpenoids | Sesquiterpenoids | Count |
| Terpenoids | Steroids | Count |
| Terpenoids | Triterpenoids | Count |
| Alkaloids | Chromanes | Count |
| Alkaloids | Diterpenoids | Count |
| Alkaloids | Fatty Acids and Conjugates | Count |
| Alkaloids | Fatty acyls | Count |
| Alkaloids | Linear polyketides | Count |
| Alkaloids | Lysine alkaloids | Count |
| Alkaloids | Macrolides | Count |
| Alkaloids | Mitomycin derivatives | Count |
| Alkaloids | Phenolic acids C6-C1 | Count |
| Alkaloids | Polyols | Count |
| Alkaloids | Pseudoalkaloids transamidation | Count |
| Alkaloids | Sphingolipids | Count |
| Alkaloids | Steroids | Count |
| Alkaloids | Triterpenoids | Count |
| Amino acids and Peptides | Anthranilic acid alkaloids | Count |
| Amino acids and Peptides | Coumarins | Count |
| Amino acids and Peptides | Fatty amides | Count |
| Amino acids and Peptides | Linear polyketides | Count |
| Amino acids and Peptides | Lysine alkaloids | Count |
| Amino acids and Peptides | Ornithine alkaloids | Count |
| Amino acids and Peptides | Pseudoalkaloids | Count |
| Amino acids and Peptides | Steroids | Count |
| Carbohydrates | Fatty esters | Count |
| Carbohydrates | Pseudoalkaloids | Count |
| Fatty acids | Lysine alkaloids | Count |
| Fatty acids | Nucleosides | Count |
| Fatty acids | Oligopeptides | Count |
| Fatty acids | Ornithine alkaloids | Count |
| Fatty acids | Phenylpropanoids C6-C3 | Count |
| Fatty acids | Pseudoalkaloids transamidation | Count |
| Fatty acids | Pseudoalkaloids | Count |
| Polyketides | Chromanes | Count |
| Polyketides | Coumarins | Count |
| Polyketides | Diphenyl ethers DPEs | Count |
| Polyketides | Diterpenoids | Count |
| Polyketides | Fatty Acids and Conjugates | Count |
| Polyketides | Fatty amides | Count |
| Polyketides | Glycerophospholipids | Count |
| Polyketides | Lysine alkaloids | Count |
| Polyketides | Mitomycin derivatives | Count |
| Polyketides | Monoterpenoids | Count |
| Polyketides | Nucleosides | Count |
| Polyketides | Ornithine alkaloids | Count |
| Polyketides | Peptide alkaloids | Count |
| Polyketides | Proline alkaloids | Count |
| Polyketides | Small peptides | Count |
| Polyketides | Sphingolipids | Count |
| Polyketides | Triterpenoids | Count |
| Polyketides | Tryptophan alkaloids | Count |
| Shikimates and Phenylpropanoids | Anthranilic acid alkaloids | Count |
| Shikimates and Phenylpropanoids | Cyclic polyketides | Count |
| Shikimates and Phenylpropanoids | Diphenyl ethers DPEs | Count |
| Shikimates and Phenylpropanoids | Fatty acyls | Count |
| Shikimates and Phenylpropanoids | Lysine alkaloids | Count |
| Shikimates and Phenylpropanoids | Meroterpenoids | Count |
| Shikimates and Phenylpropanoids | Nicotinic acid alkaloids | Count |
| Shikimates and Phenylpropanoids | Pseudoalkaloids | Count |
| Shikimates and Phenylpropanoids | Terphenyls | Count |
| Shikimates and Phenylpropanoids | Tryptophan alkaloids | Count |
| Shikimates and Phenylpropanoids | Tyrosine alkaloids | Count |
| Terpenoids | Alkylresorcinols | Count |
| Terpenoids | Carotenoids C50 | Count |
| Terpenoids | Cyclic polyketides | Count |
| Terpenoids | Fatty Acids and Conjugates | Count |
| Terpenoids | Fatty acyls | Count |
| Terpenoids | Linear polyketides | Count |
| Terpenoids | Lysine alkaloids | Count |
| Terpenoids | Oligopeptides | Count |
| Terpenoids | Ornithine alkaloids | Count |
| Terpenoids | Pseudoalkaloids transamidation | Count |
| Terpenoids | Pseudoalkaloids | Count |
| Terpenoids | Sesterterpenoids | Count |
| Terpenoids | Small peptides | Count |
| Terpenoids | Sphingolipids | Count |
| Terpenoids | Tryptophan alkaloids | Count |
| Alkaloids | Coumarins | Count |
| Alkaloids | Glycerophospholipids | Count |
| Amino acids and Peptides | Cyclic polyketides | Count |
| Amino acids and Peptides | Histidine alkaloids | Count |
| Amino acids and Peptides | Phenolic acids C6-C1 | Count |
| Amino acids and Peptides | Polycyclic aromatic polyketides | Count |
| Amino acids and Peptides | Polyols | Count |
| Carbohydrates | Amino acid glycosides | Count |
| Carbohydrates | Apocarotenoids | Count |
| Carbohydrates | Nicotinic acid alkaloids | Count |
| Carbohydrates | Phenolic acids C6-C1 | Count |
| Carbohydrates | Phloroglucinols | Count |
| Carbohydrates | Tryptophan alkaloids | Count |
| Carbohydrates | Tyrosine alkaloids | Count |
| Fatty acids | Octadecanoids | Count |
| Polyketides | Fatty acyl glycosides | Count |
| Polyketides | Flavonoids | Count |
| Polyketides | Phenolic acids C6-C1 | Count |
| Polyketides | Sesquiterpenoids | Count |
| Polyketides | Tyrosine alkaloids | Count |
| Shikimates and Phenylpropanoids | Aromatic polyketides | Count |
| Shikimates and Phenylpropanoids | Diarylheptanoids | Count |
| Shikimates and Phenylpropanoids | Diazotetronic acids and derivatives | Count |
| Shikimates and Phenylpropanoids | Fatty Acids and Conjugates | Count |
| Shikimates and Phenylpropanoids | Fluorenes | Count |
| Shikimates and Phenylpropanoids | Phenanthrenoids | Count |
| Shikimates and Phenylpropanoids | Phenylethanoids C6-C2 | Count |
| Shikimates and Phenylpropanoids | Saccharides | Count |
| Terpenoids | Aromatic polyketides | Count |
| Terpenoids | Coumarins | Count |
| Terpenoids | Eicosanoids | Count |
| Terpenoids | Fatty acyl glycosides | Count |
| Terpenoids | Flavonoids | Count |
| Terpenoids | Glycerolipids | Count |
| Terpenoids | Mycosporine derivatives | Count |
| Terpenoids | Octadecanoids | Count |
| Terpenoids | Phenolic acids C6-C1 | Count |
| Terpenoids | Polyprenols | Count |
| Terpenoids | Saccharides | Count |
| Terpenoids | Stilbenoids | Count |

**Table S2c.** Molecular traits represented by 241 molecular descriptors. The first column contains the high-level information summarizing on mathematically similar molecular descriptors describing similar chemical functions. The second column contains the low-level information represented by the respective identifiers of the CDK nomenclature. The last column contains a description of the molecular trait.

| **Molecular descriptor trait (High-level)** | **Molecular descriptor trait (Low-level)** | **Type** | **Description** |
| --- | --- | --- | --- |
| Molecular Mass | MolWeight | Numeric | Total molecular mass |
| Atom Count Descriptor | nAtoms | Ordinal | Number of atoms |
| Number of C atoms | numC | Ordinal | Sum of C atoms |
| Number of N atoms | numN | Ordinal | Sum of N atoms |
| Number of P atoms | numP | Ordinal | Sum of P atoms |
| Number of O atoms | numO | Ordinal | Sum of O atoms |
| Number of Hydrogen atoms | numHydrogen | Ordinal | Sum of Hydrogen atoms |
| CNRatio | CNRatio | Numeric | C:N ratio of molecules |
| Partition Descriptor | XLogP | Numeric | Partition coefficient represented by XLogP |
| Molecular Non-flatness Descriptor | Fsp3 | Numeric | Fractional CSP3 descriptor characterizing non-flatness of a molecule |
| Molecular Weight | MW | Numeric | Total molecular weight |
| Drug Likeness | LipinskiFailures | Numeric | Drug likeness expressed by rule of five descriptor |
| Molecular Bond Descriptor | nRotB | Numeric | Rotatable Bonds Count Descriptor |
| Partition Descriptor | MLogP | Numeric | Partition coefficient represented Mannhold LogP |
| Atom Count Descriptor | nAtomLAC | Ordinal | Number of atoms in the longest aliphatic chain |
| Atom Count Descriptor | nAtomP | Ordinal | Number of P atoms in aliphatic chain |
| Atom Count Descriptor | nAtomLC | Ordinal | Number of atoms in longest chain |
| Molecular Bond Descriptor | nB | Ordinal | Number of H bonds |
| Basic Group Count Descriptor | nBase | Ordinal | Number of basic groups |
| Atom Count Descriptor | nAtom | Ordinal | Number of atoms |
| Aromaticity Descriptor | nAromBond | Ordinal | Number of aromatic bonds |
| Aromaticity Descriptor | naAromAtom | Ordinal | Number of aromatic atoms |
| Partition Descriptor | ALogP | Numeric | Partition coefficient represented by ALogP |
| Partition Descriptor | ALogp2 | Numeric | Partition coefficient represented by ALogP^2^ |
| Partition Descriptor | AMR | Numeric | Partition coefficient represented by ALogP Molar Refractivity |
| Acidity Descriptor | nAcid | Ordinal | Acidic Group Counts |
| Amino Acid Count Descriptor | nA | Ordinal | Alanine Group Count |
| Amino Acid Count Descriptor | nR | Ordinal | Arginine Group Count |
| Amino Acid Count Descriptor | nN | Ordinal | Asparagine Group Count |
| Amino Acid Count Descriptor | nD | Ordinal | Aspartic acid Group Count |
| Amino Acid Count Descriptor | nC | Ordinal | Cysteine Group Count |
| Amino Acid Count Descriptor | nF | Ordinal | Phenylalanine Group Count |
| Amino Acid Count Descriptor | nQ | Ordinal | Glutamine Group Count |
| Amino Acid Count Descriptor | nE | Ordinal | Glutamic acid Group Count |
| Amino Acid Count Descriptor | nG | Ordinal | Glycine Group Count |
| Amino Acid Count Descriptor | nH | Ordinal | Histidine Group Count |
| Amino Acid Count Descriptor | nI | Ordinal | Isoleucine Group Count |
| Amino Acid Count Descriptor | nP | Ordinal | Proline Group Count |
| Amino Acid Count Descriptor | nL | Ordinal | Leucine Group Count |
| Amino Acid Count Descriptor | nK | Ordinal | Lysine Group Count |
| Amino Acid Count Descriptor | nM | Ordinal | Methionine Group Count |
| Amino Acid Count Descriptor | nS | Ordinal | Serine Group Count |
| Amino Acid Count Descriptor | nT | Ordinal | Threonine Group Count |
| Amino Acid Count Descriptor | nY | Ordinal | Tyrosine Group Count |
| Amino Acid Count Descriptor | nV | Ordinal | Valine Group Count |
| Amino Acid Count Descriptor | nW | Ordinal | Tryptophan Group Count |
| Molecular Electronic Descriptor | apol | Numeric | The sum of the atomic polarizabilities (including implicit hydrogens) |
| Molecular Electronic Descriptor | bpol | Numeric | the sum of the absolute value of the difference between atomic polarizabilities of all bonded atoms in the molecule (including implicit hydrogens) |
| Molecular Ring Descriptor | nSmallRings | Ordinal | Total number of small rings |
| Molecular Ring Descriptor | nAromRings | Ordinal | Total number of small aromatic rings |
| Molecular Ring Descriptor | nRingBlocks | Ordinal | Total number of distinct ring blocks |
| Molecular Ring Descriptor | nAromBlocks | Ordinal | Total number of aromatically connected components |
| Molecular Ring Descriptor | nRings3 | Numeric | Individual breakdown of 3 membered rings |
| Molecular Ring Descriptor | nRings4 | Numeric | Individual breakdown of 4 membered rings |
| Molecular Ring Descriptor | nRings5 | Numeric | Individual breakdown of 5 membered rings |
| Molecular Ring Descriptor | nRings6 | Numeric | Individual breakdown of 6 membered rings |
| Molecular Ring Descriptor | nRings7 | Numeric | Individual breakdown of 7 membered rings |
| Molecular Ring Descriptor | nRings8 | Numeric | Individual breakdown of 8 membered rings |
| Molecular Ring Descriptor | nRings9 | Numeric | Individual breakdown of 9 membered rings |
| Molecular Topological Descriptor | tpsaEfficiency | Numeric | Topological polar surface area which reflects the ability of a molecule to penetrate into a cell |
| Eccentric Connectivity Descriptor | Zagreb | Numeric | Eccentric Connectivity expressed through Zagreb Index Descriptor |
| Molecular Path Descriptor | WPATH | Numeric | Wiener path number |
| Molecular Polarity Descriptor | WPOL | Numeric | Wiener polarity number |
| Molecular Path Descriptor | WTPT.1 | Numeric | The weighted path (1) descriptor described by Randic characterizing molecular branching |
| Molecular Path Descriptor | WTPT.2 | Numeric | The weighted path (2) descriptor described by Randic characterizing molecular branching |
| Molecular Path Descriptor | WTPT.3 | Numeric | The weighted path (3) descriptor described by Randic characterizing molecular branching |
| Molecular Path Descriptor | WTPT.4 | Numeric | The weighted path (4) descriptor described by Randic characterizing molecular branching |
| Molecular Path Descriptor | WTPT.5 | Numeric | The weighted path (5) descriptor described by Randic characterizing molecular branching |
| Molecular Topological Descriptor | VAdjMat | Numeric | Vertex adjacency information of a molecule |
| Molecular Topological Descriptor | VABC | Numeric | Van der Waals Volume |
| Molecular Topological Descriptor | TopoPSA | Numeric | Topological polar surface area |
| Molecular Topological Descriptor | topoShape | Numeric | Petitjean topological shape index describing the anisotropy in a molecule |
| Molecular Topological Descriptor | geomShape | Numeric | Petitjean topological shape index describing the anisotropy in a molecule |
| Molecular Topological Descriptor | PetitjeanNumber | Numeric | Petitjean Number of a molecule |
| Molecular Distance Edge Descriptor | MDEC.11 | Numeric | Molecular distance edge descriptors for C, N and O |
| Molecular Distance Edge Descriptor | MDEC.12 | Numeric | Molecular distance edge descriptors for C, N and O |
| Molecular Distance Edge Descriptor | MDEC.13 | Numeric | Molecular distance edge descriptors for C, N and O |
| Molecular Distance Edge Descriptor | MDEC.14 | Numeric | Molecular distance edge descriptors for C, N and O |
| Molecular Distance Edge Descriptor | MDEC.22 | Numeric | Molecular distance edge descriptors for C, N and O |
| Molecular Distance Edge Descriptor | MDEC.23 | Numeric | Molecular distance edge descriptors for C, N and O |
| Molecular Distance Edge Descriptor | MDEC.24 | Numeric | Molecular distance edge descriptors for C, N and O |
| Molecular Distance Edge Descriptor | MDEC.33 | Numeric | Molecular distance edge descriptors for C, N and O |
| Molecular Distance Edge Descriptor | MDEC.34 | Numeric | Molecular distance edge descriptors for C, N and O |
| Molecular Distance Edge Descriptor | MDEC.44 | Numeric | Molecular distance edge descriptors for C, N and O |
| Molecular Distance Edge Descriptor | MDEO.11 | Numeric | Molecular distance edge descriptors for C, N and O |
| Molecular Distance Edge Descriptor | MDEO.12 | Numeric | Molecular distance edge descriptors for C, N and O |
| Molecular Distance Edge Descriptor | MDEO.22 | Numeric | Molecular distance edge descriptors for C, N and O |
| Molecular Distance Edge Descriptor | MDEN.11 | Numeric | Molecular distance edge descriptors for C, N and O |
| Molecular Distance Edge Descriptor | MDEN.12 | Numeric | Molecular distance edge descriptors for C, N and O |
| Molecular Distance Edge Descriptor | MDEN.13 | Numeric | Molecular distance edge descriptors for C, N and O |
| Molecular Distance Edge Descriptor | MDEN.22 | Numeric | Molecular distance edge descriptors for C, N and O |
| Molecular Distance Edge Descriptor | MDEN.23 | Numeric | Molecular distance edge descriptors for C, N and O |
| Molecular Distance Edge Descriptor | MDEN.33 | Numeric | Molecular distance edge descriptors for C, N and O |
| Molecular Electron State Descriptor | khs.sLi | Numeric | Electron state for chemical group -Li |
| Molecular Electron State Descriptor | khs.ssBe | Numeric | Electron state for chemical group -Be- |
| Molecular Electron State Descriptor | khs.ssssBe | Numeric | Electron state for chemical group >Be<^-2^ |
| Molecular Electron State Descriptor | khs.ssBH | Numeric | Electron state for chemical group -BH- |
| Molecular Electron State Descriptor | khs.sssB | Numeric | Electron state for chemical group -B< |
| Molecular Electron State Descriptor | khs.ssssB | Numeric | Electron state for chemical group >B<- |
| Molecular Electron State Descriptor | khs.sCH3 | Numeric | Electron state for chemical group -CH_3_ |
| Molecular Electron State Descriptor | khs.dCH2 | Numeric | Electron state for chemical group =CH_2_ |
| Molecular Electron State Descriptor | khs.ssCH2 | Numeric | Electron state for chemical group -CH^2-^ |
| Molecular Electron State Descriptor | khs.tCH | Numeric | Electron state for chemical group #CH |
| Molecular Electron State Descriptor | khs.dsCH | Numeric | Electron state for chemical group =CH- |
| Molecular Electron State Descriptor | khs.aaCH | Numeric | Electron state for chemical group aCHa |
| Molecular Electron State Descriptor | khs.sssCH | Numeric | Electron state for chemical group >CH- |
| Molecular Electron State Descriptor | khs.ddC | Numeric | Electron state for chemical group =C= |
| Molecular Electron State Descriptor | khs.tsC | Numeric | Electron state for chemical group #C- |
| Molecular Electron State Descriptor | khs.dssC | Numeric | Electron state for chemical group =C< |
| Molecular Electron State Descriptor | khs.aasC | Numeric | Electron state for chemical group aCa- |
| Molecular Electron State Descriptor | khs.aaaC | Numeric | Electron state for chemical group aaCa |
| Molecular Electron State Descriptor | khs.ssssC | Numeric | Electron state for chemical group >C< |
| Molecular Electron State Descriptor | khs.sNH3 | Numeric | Electron state for chemical group -NH_3_^+^ |
| Molecular Electron State Descriptor | khs.sNH2 | Numeric | Electron state for chemical group -NH_2_ |
| Molecular Electron State Descriptor | khs.ssNH2 | Numeric | Electron state for chemical group -NH_2_-^+^ |
| Molecular Electron State Descriptor | khs.dNH | Numeric | Electron state for chemical group =NH |
| Molecular Electron State Descriptor | khs.ssNH | Numeric | Electron state for chemical group -NH- |
| Molecular Electron State Descriptor | khs.aaNH | Numeric | Electron state for chemical group aNHa |
| Molecular Electron State Descriptor | khs.tN | Numeric | Electron state for chemical group #N |
| Molecular Electron State Descriptor | khs.sssNH | Numeric | Electron state for chemical group >NH-^+^ |
| Molecular Electron State Descriptor | khs.dsN | Numeric | Electron state for chemical group =N- |
| Molecular Electron State Descriptor | khs.aaN | Numeric | Electron state for chemical group aNa |
| Molecular Electron State Descriptor | khs.sssN | Numeric | Electron state for chemical group >N- |
| Molecular Electron State Descriptor | khs.ddsN | Numeric | Electron state for chemical group -N<< |
| Molecular Electron State Descriptor | khs.aasN | Numeric | Electron state for chemical group aaNs |
| Molecular Electron State Descriptor | khs.ssssN | Numeric | Electron state for chemical group >N<^+^ |
| Molecular Electron State Descriptor | khs.sOH | Numeric | Electron state for chemical group -OH |
| Molecular Electron State Descriptor | khs.dO | Numeric | Electron state for chemical group=O |
| Molecular Electron State Descriptor | khs.ssO | Numeric | Electron state for chemical group -O- |
| Molecular Electron State Descriptor | khs.aaO | Numeric | Electron state for chemical group aOa |
| Molecular Electron State Descriptor | khs.sF | Numeric | Electron state for chemical group -F |
| Molecular Electron State Descriptor | khs.sSiH3 | Numeric | Electron state for chemical group -SiH3 |
| Molecular Electron State Descriptor | khs.ssSiH2 | Numeric | Electron state for chemical group -SiH2- |
| Molecular Electron State Descriptor | khs.sssSiH | Numeric | Electron state for chemical group >SiH- |
| Molecular Electron State Descriptor | khs.ssssSi | Numeric | Electron state for chemical group >Si< |
| Molecular Electron State Descriptor | khs.sPH2 | Numeric | Electron state for chemical group -PH2 |
| Molecular Electron State Descriptor | khs.ssPH | Numeric | Electron state for chemical group -PH- |
| Molecular Electron State Descriptor | khs.sssP | Numeric | Electron state for chemical group >P- |
| Molecular Electron State Descriptor | khs.dsssP | Numeric | Electron state for chemical group ->P= |
| Molecular Electron State Descriptor | khs.sssssP | Numeric | Electron state for chemical group ->P< |
| Molecular Electron State Descriptor | khs.sSH | Numeric | Electron state for chemical group -SH |
| Molecular Electron State Descriptor | khs.dS | Numeric | Electron state for chemical group =S |
| Molecular Electron State Descriptor | khs.ssS | Numeric | Electron state for chemical group -S- |
| Molecular Electron State Descriptor | khs.aaS | Numeric | Electron state for chemical group aSa |
| Molecular Electron State Descriptor | khs.dssS | Numeric | Electron state for chemical group >S= |
| Molecular Electron State Descriptor | khs.ddssS | Numeric | Electron state for chemical group >S== |
| Molecular Electron State Descriptor | Khs.ssssssS | Numeric | Electron state for chemical group >S<< |
| Molecular Electron State Descriptor | khs.sCl | Numeric | Electron state for chemical group -Cl |
| Molecular Electron State Descriptor | khs.sGeH3 | Numeric | Electron state for chemical group -GeH3 |
| Molecular Electron State Descriptor | khs.ssGeH2 | Numeric | Electron state for chemical group -GeH2- |
| Molecular Electron State Descriptor | khs.sssGeH | Numeric | Electron state for chemical group >GeH- |
| Molecular Electron State Descriptor | khs.ssssGe | Numeric | Electron state for chemical group >Ge< |
| Molecular Electron State Descriptor | khs.sAsH2 | Numeric | Electron state for chemical group -AsH2 |
| Molecular Electron State Descriptor | khs.ssAsH | Numeric | Electron state for chemical group -AsH- |
| Molecular Electron State Descriptor | khs.sssAs | Numeric | Electron state for chemical group >As- |
| Molecular Electron State Descriptor | khs.sssdAs | Numeric | Electron state for chemical group ->As= |
| Molecular Electron State Descriptor | khs.sssssAs | Numeric | Electron state for chemical group ->As< |
| Molecular Electron State Descriptor | khs.sSeH | Numeric | Electron state for chemical group -SeH |
| Molecular Electron State Descriptor | khs.dSe | Numeric | Electron state for chemical group =Se |
| Molecular Electron State Descriptor | khs.ssSe | Numeric | Electron state for chemical group -Se- |
| Molecular Electron State Descriptor | khs.aaSe | Numeric | Electron state for chemical group aSea |
| Molecular Electron State Descriptor | khs.dssSe | Numeric | Electron state for chemical group >Se= |
| Molecular Electron State Descriptor | khs.ddssSe | Numeric | Electron state for chemical group =>Se< |
| Molecular Electron State Descriptor | khs.sBr | Numeric | Electron state for chemical group -Br |
| Molecular Electron State Descriptor | khs.sSnH3 | Numeric | Electron state for chemical group -SnH3 |
| Molecular Electron State Descriptor | khs.ssSnH2 | Numeric | Electron state for chemical group -SnH2- |
| Molecular Electron State Descriptor | khs.sssSnH | Numeric | Electron state for chemical group >SnH- |
| Molecular Electron State Descriptor | khs.ssssSn | Numeric | Electron state for chemical group >Sn< |
| Molecular Electron State Descriptor | khs.sI | Numeric | Electron state for chemical group -I |
| Molecular Electron State Descriptor | khs.sPbH3 | Numeric | Electron state for chemical group -PbH3 |
| Molecular Electron State Descriptor | khs.ssPbH2 | Numeric | Electron state for chemical group -PbH2- |
| Molecular Electron State Descriptor | khs.sssPbH | Numeric | Electron state for chemical group >PbH- |
| Molecular Electron State Descriptor | khs.ssssPb | Numeric | Electron state for chemical group >Pb< |
| Molecular Shape Descriptor | Kier1 | Numeric | Kier and Hall kappa molecular shape index |
| Molecular Shape Descriptor | Kier2 | Numeric | Kier and Hall kappa molecular shape index |
| Molecular Shape Descriptor | Kier3 | Numeric | Kier and Hall kappa molecular shape index |
| Hybridization Ratio Descriptor | HybRatio | Numeric | Hybridization Ratio |
| Molecular Complexity Descriptor | fragC | Numeric | Fragment Complexity |
| Molecular Complexity Descriptor | FMF | Numeric | FMF Descriptor characterizing complexity of a molecule |
| Eccentric Connectivity Descriptor | ECCEN | Numeric | Eccentric connectivity index |
| Molecular Topological Descriptor | SP.0 | Numeric | Kier and Hall Chi path index of order 0 |
| Molecular Topological Descriptor | SP.1 | Numeric | Kier and Hall Chi path index of order 1 |
| Molecular Topological Descriptor | SP.2 | Numeric | Kier and Hall Chi path index of order 2 |
| Molecular Topological Descriptor | SP.3 | Numeric | Kier and Hall Chi path index of order 3 |
| Molecular Topological Descriptor | SP.4 | Numeric | Kier and Hall Chi path index of order 4 |
| Molecular Topological Descriptor | SP.5 | Numeric | Kier and Hall Chi path index of order 5 |
| Molecular Topological Descriptor | SP.6 | Numeric | Kier and Hall Chi path index of order 6 |
| Molecular Topological Descriptor | SP.7 | Numeric | Kier and Hall Chi path index of order 7 |
| Molecular Topological Descriptor | VP.0 | Numeric | Kier and Hall Chi path index of order 0 |
| Molecular Topological Descriptor | VP.1 | Numeric | Kier and Hall Chi path index of order 1 |
| Molecular Topological Descriptor | VP.2 | Numeric | Kier and Hall Chi path index of order 2 |
| Molecular Topological Descriptor | VP.3 | Numeric | Kier and Hall Chi path index of order 3 |
| Molecular Topological Descriptor | VP.4 | Numeric | Kier and Hall Chi path index of order 4 |
| Molecular Topological Descriptor | VP.5 | Numeric | Kier and Hall Chi path index of order 5 |
| Molecular Topological Descriptor | VP.6 | Numeric | Kier and Hall Chi path index of order 6 |
| Molecular Topological Descriptor | VP.7 | Numeric | Kier and Hall Chi path index of order 7 |
| Molecular Topological Descriptor | SPC.4 | Numeric | Kier and Hall Chi path cluster index of order 4 |
| Molecular Topological Descriptor | SPC.5 | Numeric | Kier and Hall Chi path cluster index of order 5 |
| Molecular Topological Descriptor | SPC.6 | Numeric | Kier and Hall Chi path cluster index of order 6 |
| Molecular Topological Descriptor | VPC.4 | Numeric | Kier and Hall Chi path cluster index of order 4 |
| Molecular Topological Descriptor | VPC.5 | Numeric | Kier and Hall Chi path cluster index of order 5 |
| Molecular Topological Descriptor | VPC.6 | Numeric | Kier and Hall Chi path cluster index of order 6 |
| Molecular Topological Descriptor | SC.3 | Numeric | Kier and Hall Chi cluster index of order 3 |
| Molecular Topological Descriptor | SC.4 | Numeric | Kier and Hall Chi cluster index of order 4 |
| Molecular Topological Descriptor | SC.5 | Numeric | Kier and Hall Chi cluster index of order 5 |
| Molecular Topological Descriptor | SC.6 | Numeric | Kier and Hall Chi cluster index of order 6 |
| Molecular Topological Descriptor | VC.3 | Numeric | Kier and Hall Chi cluster index of order 3 |
| Molecular Topological Descriptor | VC.4 | Numeric | Kier and Hall Chi cluster index of order 4 |
| Molecular Topological Descriptor | VC.5 | Numeric | Kier and Hall Chi cluster index of order 5 |
| Molecular Topological Descriptor | VC.6 | Numeric | Kier and Hall Chi cluster index of order 6 |
| Molecular Topological Descriptor | SCH.3 | Numeric | Kier and Hall Chi chain index of order 3 |
| Molecular Topological Descriptor | SCH.4 | Numeric | Kier and Hall Chi chain index of order 4 |
| Molecular Topological Descriptor | SCH.5 | Numeric | Kier and Hall Chi chain index of order 5 |
| Molecular Topological Descriptor | SCH.6 | Numeric | Kier and Hall Chi chain index of order 6 |
| Molecular Topological Descriptor | SCH.7 | Numeric | Kier and Hall Chi chain index of order 7 |
| Molecular Topological Descriptor | VCH.3 | Numeric | Kier and Hall Chi chain index of order 3 |
| Molecular Topological Descriptor | VCH.4 | Numeric | Kier and Hall Chi chain index of order 4 |
| Molecular Topological Descriptor | VCH.5 | Numeric | Kier and Hall Chi chain index of order 5 |
| Molecular Topological Descriptor | VCH.6 | Numeric | Kier and Hall Chi chain index of order 6 |
| Molecular Topological Descriptor | VCH.7 | Numeric | Kier and Hall Chi chain index of order 7 |
| Carbon Connectivity Descriptor | C1SP1 | Numeric | Carbon connectivity in terms of hybridization |
| Carbon Connectivity Descriptor | C2SP1 | Numeric | Carbon connectivity in terms of hybridization |
| Carbon Connectivity Descriptor | C1SP2 | Numeric | Carbon connectivity in terms of hybridization |
| Carbon Connectivity Descriptor | C2SP2 | Numeric | Carbon connectivity in terms of hybridization |
| Carbon Connectivity Descriptor | C3SP2 | Numeric | Carbon connectivity in terms of hybridization |
| Carbon Connectivity Descriptor | C1SP3 | Numeric | Carbon connectivity in terms of hybridization |
| Carbon Connectivity Descriptor | C2SP3 | Numeric | Carbon connectivity in terms of hybridization |
| Carbon Connectivity Descriptor | C3SP3 | Numeric | Carbon connectivity in terms of hybridization |
| Carbon Connectivity Descriptor | C4SP3 | Numeric | Carbon connectivity in terms of hybridization |
| Molecular Topological Descriptor | ATSp1 | Numeric | The Moreau-Broto autocorrelation descriptor using polarizability |
| Molecular Topological Descriptor | ATSp2 | Numeric | The Moreau-Broto autocorrelation descriptor using polarizability |
| Molecular Topological Descriptor | ATSp3 | Numeric | The Moreau-Broto autocorrelation descriptor using polarizability |
| Molecular Topological Descriptor | ATSp4 | Numeric | The Moreau-Broto autocorrelation descriptor using polarizability |
| Molecular Topological Descriptor | ATSp5 | Numeric | The Moreau-Broto autocorrelation descriptor using polarizability |
| Molecular Topological Descriptor | ATSm1 | Numeric | The Moreau-Broto autocorrelation descriptor using atomic weight |
| Molecular Topological Descriptor | ATSm2 | Numeric | The Moreau-Broto autocorrelation descriptor using atomic weight |
| Molecular Topological Descriptor | ATSm3 | Numeric | The Moreau-Broto autocorrelation descriptor using atomic weight |
| Molecular Topological Descriptor | ATSm4 | Numeric | The Moreau-Broto autocorrelation descriptor using atomic weight |
| Molecular Topological Descriptor | ATSm5 | Numeric | The Moreau-Broto autocorrelation descriptor using atomic weight |
| Molecular Topological Descriptor | ATSc1 | Numeric | The Moreau-Broto autocorrelation descriptor using partial charges |
| Molecular Topological Descriptor | ATSc2 | Numeric | The Moreau-Broto autocorrelation descriptor using partial charges |
| Molecular Topological Descriptor | ATSc3 | Numeric | The Moreau-Broto autocorrelation descriptor using partial charges |
| Molecular Topological Descriptor | ATSc4 | Numeric | The Moreau-Broto autocorrelation descriptor using partial charges |
| Molecular Topological Descriptor | ATSc5 | Numeric | The Moreau-Broto autocorrelation descriptor using partial charges |
